# Genetically Programmed Shape-morphing of Engineered Living Materials

**DOI:** 10.64898/2026.03.05.709855

**Authors:** Jan Becker, Yuchen Liu, Miguel Baños, Rosanne Schmachtenberg, Mahmudul Hasan, Claudia Fink-Straube, Luai R. Khoury, Wilfried Weber

## Abstract

Engineered living materials (ELMs) promise genetically programmable functions by coupling biological regulation to synthetic material responses. Here, we introduce genetically encoded, reversible shape-morphing in a peptide-crosslinked polyethylene glycol (PEG) hydrogel whose network density is modulated by opposing enzymatic pairs that induce crosslinking or hydrolysis. This molecular programmability alternates the hydrogel between deswelling and swelling/disintegration and produces 2 - 5-fold changes in mechanical properties. By fabricating a bilayer hydrogel with an inert layer, these molecular modulations are translated into a reversible and directional motion with angular bending motions exceeding 80°. Further, by embedding genetically engineered bacteria or interfacing mammalian cells, producing the relevant enzymatic cues, the reversible shape-morphing of these ELMs is programmed at the genetic level. We further demonstrate genetically programmed, autonomous reversible bending in a bilayer hydrogel controlled by out-of-equilibrium counteracting biochemical reactions with dynamically changing respective reaction rates. This work establishes a concept where coordinated polymer/peptide material engineering and synthetic biology yield autonomous shape-morphing ELMs, opening avenues toward biohybrid soft robotics, adaptive microfluidic systems, and dynamic biomedical interfaces.

## 1. Introduction

Engineered living materials (ELMs) represent a rapidly emerging class of materials in which living cells contribute advanced properties and functions such as adaptation to external cues. These biohybrid systems couple molecular sensing, genetic regulation, and biological actuation to convert multimodal cues into adaptive material responses, yielding adaptivity far beyond conventional stimuli-responsive approaches.^[1–5]^ This enables real-time, programmable functions such as targeted release, self-healing, selective degradation, and controlled stiffening within sustainable, biocompatible platforms for advanced biomaterials, therapeutics, and biosensors.^[1–8]^

Hydrogels serve as ideal material for ELM synthesis due to their adjustable biochemical functionality, tunable physical properties and excellent biocompatibility. Their hydrated 3D polymer networks mimic soft tissues, highlighting their suitability as a bio-programmable matrix.^[9,10]^ The modularity provided by these systems allows tailoring the polymer backbone and crosslinking mechanism to enable diverse applications in regenerative medicine^[9–15]^, soft devices^[16,17]^, microfluidics^[16–19]^, drug delivery^[20–22]^, and bioremediation^[23–25]^.

Recently, shape-morphing hydrogels have gained prominence by translating environmental cues into programmed mechanical deformations such as bending, twisting or buckling, thereby imitating movements found in nature. Such deformations can be achieved by adjusting crosslinking density, swelling ratio, stiffness, osmolarity, and polymer (chain/fiber) alignment, enabling local molecular-scale changes to be amplified into programmed macroscopic actuation, thereby unlocking functionalities that are inaccessible to static networks.^[26,27]^ Spatial patterning during the fabrication process via photomasking, layered architectures, or 3D-printing enables the design of locally confined actuation regions, which in turn supports controlled, directional movement and complex shape trajectories.^[28–31]^ Such shape transformations have been exploited across a broad application space, including reversible gripping and release in soft robotic manipulators^[16,28,32,33]^, curvature generation and lumen formation in tissue morphogenesis models^[34,35]^, stimulus-gated valving and flow regulation in microfluidic systems^[36,37]^, and on-demand exposure or concealment of therapeutic payloads for spatiotemporally controlled delivery^[14,38]^.

Shape-morphing actuation can be initiated by chemical (e.g., pH^[28,39]^, ionic strength^[31,40,41]^), physical (e.g., temperature^[29,30,42]^, light^[43]^, magnetic fields^[44,45]^), or biological (e.g., enzymes^[38,46–48]^ or cell proliferation^[49,50]^) stimuli, each offering distinct advantages and constraints.^[26,27]^ Reversible physicochemical triggers (pH/ionic strength and temperature) and externally applied fields (e.g., light, magnetic) are often well-suited for repeatable cycling and rapid switching. However, they can compromise spatial precision (particularly for temperature in bulk environments), exhibit threshold-like “all-or-none” responses with limited tunability around narrow operating windows, or proceed too rapidly for applications requiring sustained, controlled release.^[16,27,48,51]^ By contrast, covalent bond-forming or -breaking reactions (such as crosslinking and hydrolysis) offer a more stable, but typically less repeatable, route to deformation, while providing higher mechanical robustness and retention of the programmed shape even after the initiating cue has faded.^[26,27,29,38,40,52]^

Enzymes have shown great potential as stimuli for responsive biomaterials^[53–58]^ and are currently being explored to induce shape-morphing in biological systems.^[47,48,51]^ By integrating peptide- or protein-based substrates within the material, enzymes can be used to remodel the polymer networks by breaking or forming (iso-)peptide crosslinks, thereby modulating physical material properties.^[38,52,57,59]^ Further, enzymatic activity can be spatially confined by means of enzyme diffusion potential and localization, as well as substrate sequence distribution, enabling targeted material modifications.^[60,61]^ In addition, enzymes provide an extensive library of actuators distinguished by their high specificity, catalytic amplification potential and ability to operate under a variety of physiological conditions, thereby mitigating cytotoxicity risks posed by stimuli like UV/blue light or extreme pH. Crucially, by genetically engineering organisms to produce the relevant enzymatic cues, hydrogel shape-morphing can be programmed at the genetic level, allowing cells to autonomously direct and actuate the process.^[1,5,6,50,62]^

In this study, we introduce the concept of programming shape-morphing into ELMs via genetically encoded enzymatic actuators. To this end, we employ short peptides containing enzyme substrate motifs to form a biologically responsive PEG-based hydrogel. We demonstrate the modulability of these materials by means of two opposing actuator pairs that facilitate enzyme-mediated crosslinking or hydrolysis of the hydrogel network. While tyrosinase or transglutaminase (TG) mediate an increase in network crosslinking density, subsequent exposure to the proteases trypsin or Pro-Pro endopeptidase-1 (PPEP-1) leads to matrix bond hydrolysis. These network modifications drive sequential and reversible deswelling/swelling with pronounced changes of mechanical properties, as confirmed by rheology. By laminating the responsive hydrogel to an inert layer, we construct a bilayer architecture that spatially confines remodeling and converts these molecular modifications into directional 3D bending. Finally, by incorporating bacteria or mammalian cells genetically engineered to produce the relevant enzymes, we synthesize ELMs that autonomously perform reversible shape morphing via counteracting biochemical reactions with dynamically changing relative reaction rates, thereby eliminating the need for external physical instruments. This genetically actuated strategy provides a route to deployable biohybrid actuators, overcoming current limitations of portability and integration, and advancing present systems towards autonomous soft robotics, self-regulating microfluidics, adaptive therapeutic implants, and smart drug delivery systems.

## 2. Results

### 2.1. Design and functionality of the biologically responsive and inert hydrogels

Towards translating enzyme-mediated molecular changes into a mechanical material response, a biologically responsive hydrogel that undergoes volumetric deswelling or swelling when exposed to the relevant actuators is required. Since elevating planar motions into a directional 3D shape-morphing relies on an asymmetric material response, we laminated inert and responsive hydrogels to fabricate bilayer constructs with spatially confined enzymatic responsivity. Upon treatment with crosslinking enzymes, additional peptide bonds were introduced exclusively into the responsive layer, inducing locally confined deswelling, increasing mechanical properties, and thereby bending of the bilayer material. Subsequent exposure to hydrolytic enzymes leads to localized peptide bond cleavage, causing swelling and decreasing of mechanical properties, hence inducing shape recovery via the elastic restoring force of the inert layer (**Figure 1A**). Beyond the initiation of deformation by manual enzyme addition, bacteria or mammalian cells can be genetically programmed to produce the required enzymatic actuators and subsequently incorporated into the materials, thereby resulting in an autonomous shape-morphing ELM.

**Figure 1.**
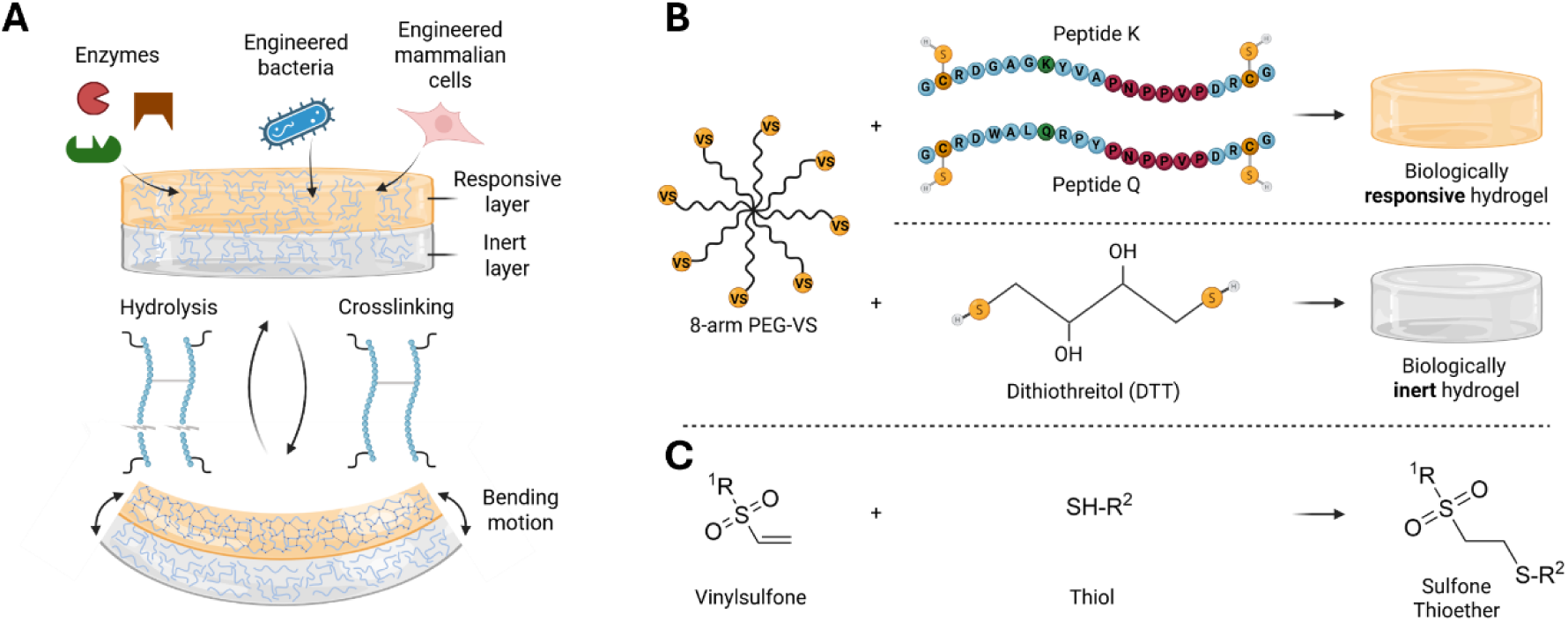
Enzyme-mediated bending mechanism and hydrogel design. (A) Enzyme-mediated bending mechanism of bilayer hydrogels. A responsive hydrogel is covalently laminated to an inert layer to generate a bilayer material with spatially restricted network remodeling. Localized crosslinking by (genetically encoded) enzymes induces one-sided deswelling and directional bending, while enzyme-mediated peptide-bond hydrolysis restores the original shape. (B) Responsive and inert hydrogel formation. Eight-arm polyethylene glycol (PEG) functionalized with vinyl sulfone (VS) forms biologically responsive hydrogels when crosslinked by peptide cysteine residues (C), or biologically inert hydrogels when crosslinked by dithiothreitol (DTT). Peptide crosslinkers were designed to contain specific motifs, enabling enzyme-driven modulation of the hydrogel network. (C) Reaction mechanism of hydrogel formation. VS group of PEG reacts with thiol group of C or DTT to form a sulfone thioether bond.

To achieve asymmetric material behavior, we designed two hydrogel types: a biologically responsive hydrogel and an inert counterpart, which share basic polymerization mechanism yet differ in swelling behavior upon enzymatic exposure. To this end, we employed peptides with cysteine residues (C) at either end, or dithiothreitol (DTT), to crosslink 8-arm PEG functionalized with vinyl sulfone (PEG-VS, Figure 1B). Crosslinking proceeded via thioether bond formation by a Michael-type addition (Figure 1C). The peptides incorporated substrate sites for subsequent modulation of network density through enzyme-mediated formation of crosslinks or hydrolysis of peptide bonds.

### 2.2. Tyrosinase and trypsin-mediated hydrogel modulation

In this study, we achieve dynamic swelling changes by adjusting network density through orthogonal enzyme pairs with opposing remodeling mechanism: the formation or the hydrolysis of crosslinks. First, we evaluated tyrosinase and trypsin as hydrogel-modulating enzymes.

#### 2.2.1. Tyrosinase and trypsin-mediated reversible swelling and mechanical properties changes

To enable tyrosinase-mediated hydrogel crosslinking, a peptide (Peptide-K) harboring a lysine (K) and a tyrosine (Y) residue was used to crosslink 8-arm PEG-VS to form the initial hydrogel matrix. While Y serves as tyrosinase substrate and allows direct crosslinking with amino groups contained in K (**Figure S1A**), K residues can further be crosslinked by tyrosinase-based oxidation of caffeic acid (CA, Figure S1B). In both cases, tyrosinase catalyses the formation of intermediate, highly reactive o-quinones which subsequently form covalent crosslinks with nucleophilic amino acid side chains or undergo quinone-quinone coupling (**Figure 2A)**. As a result, hydrogel network crosslinking density is increased causing the material to undergo volumetric deswelling.

**Figure 2.**
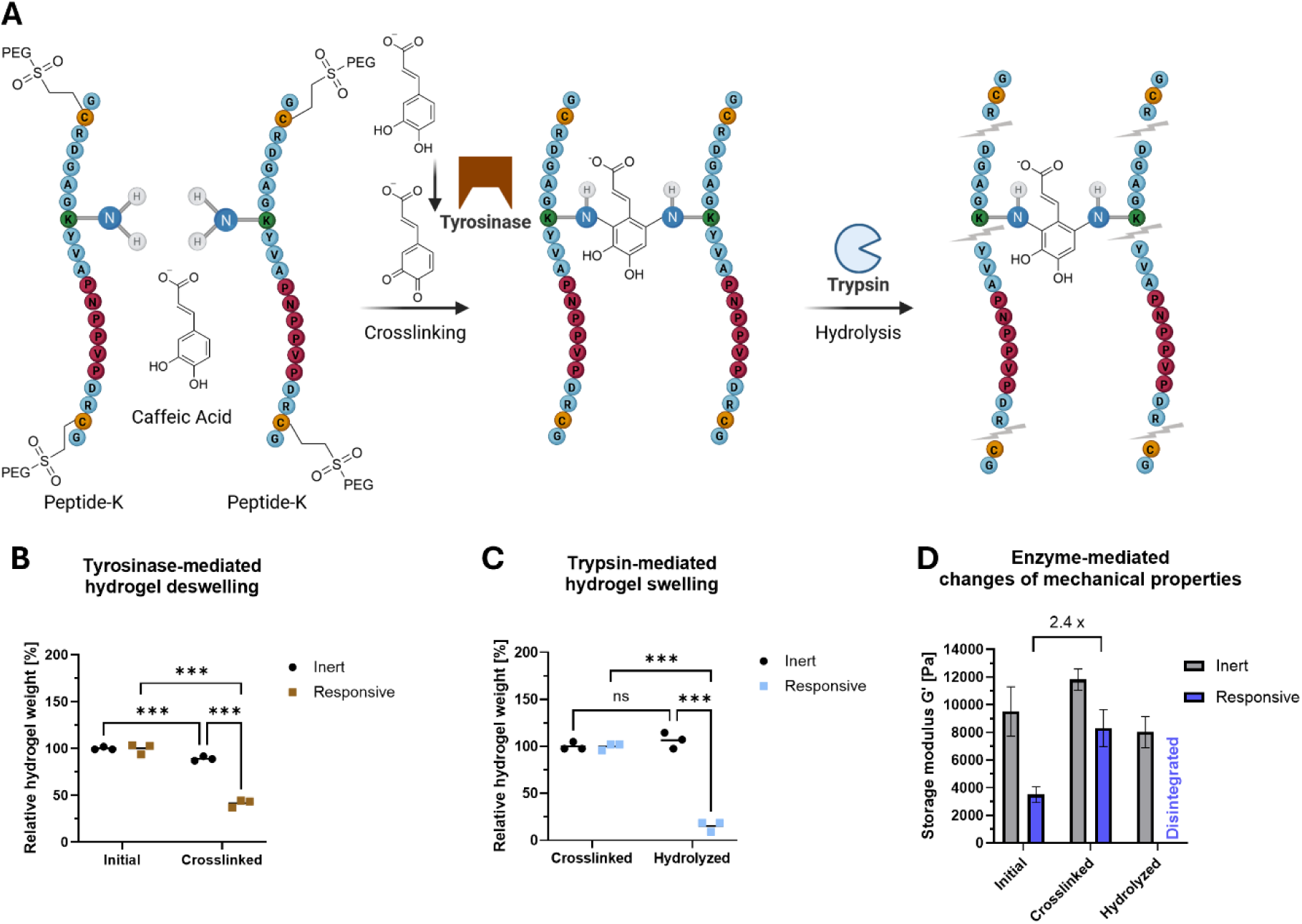
Mechanical and physical characterization of tyrosinase- and trypsin-responsive hydrogels. (A) Schematic representation of enzyme-mediated network modulation. Tyrosinase-mediated crosslinking via oxidation of caffeic acid (CA) between two lysine (K) residues contained in the peptide (Peptide-K) used to form a hydrogel with 8-arm PEG-VS. Upon addition of the protease trypsin, peptide bonds at the C-terminus of arginine (R) or K residues are hydrolyzed, fragmenting the peptide linkers and disintegrating the hydrogel. (B) Tyrosinase-mediated deswelling of hydrogels. Hydrogels were prepared as depicted in Figure 1B: 8-arm PEG-VS was crosslinked with either DTT to form an inert hydrogel, or Peptide-K to form a biologically responsive hydrogel. After 60 min polymerization time, hydrogels were equilibrated in minimal medium (M9) overnight and subsequently initial weight was recorded. Afterward, crosslinking was initiated by addition of tyrosinase (27.5 U L^-1^) and CA (5 mM) and incubated for 48 h at 37 °C before final mass was recorded. (C) Trypsin-mediated disintegration of hydrogels. Crosslinked hydrogels from (B) were washed and subsequently incubated in trypsin (5000 U L^-1^) at 37 °C and further weight changes recorded after 16 h. (D) Enzyme-mediated changes in mechanical properties of hydrogels. Storage moduli (G’) of hydrogels from (B) and (C) were recorded by small amplitude oscillatory shear (SAOS) measurements before and after enzymatic treatments. Values are shown as mean ± SD (n = 3). Normalization for hydrogel weight was performed relative to pre-treatment values, statistical significance was assessed by two-way ANOVA; ns = not significant, * = p ≤ 0.033, ** = p ≤ 0.002, *** = p ≤ 0.001.

As mechanism to reduce crosslink density and induce volumetric swelling, we evaluated the protease trypsin that cleaves peptide sequences containing arginine (R) or K residues. To this aim, we placed one R residue next to each of the two cysteine residues used to crosslink PEG-VS. Thus, trypsin addition is expected to fragment the peptides via hydrolysis of peptide bonds, disintegrating the hydrogel (Figure 2A and S1C).

To test this concept, we polymerized 8-arm PEG-VS with either Peptide-K or DTT to form responsive and inert hydrogels, respectively. Since changes in volumetric hydrogel dimension correlate with overall hydrogel weight, the resulting gels were weighed directly after polymerization and again after equilibration in minimal medium (M9, pH 6.8). This assessment of equilibration behavior revealed a minor decrease in mass for inert hydrogels to 90 ± 1% (**Figure S2A**) relative to polymerization weight while responsive hydrogels maintained their mass (101 ± 1%). Afterward, both hydrogels were treated with 27.5 tyrosinase enzyme units per liter (U L^-1^) in combination with CA (5 mM) as substrate. Notably, inert and responsive hydrogels adopted an irreversible dark-brown staining post treatment, indicating assembly of melanin-like chromophoric o-quinone polymers within the hydrogel matrix (Figure S2B).^[63]^ Determination of hydrogel weight revealed a significant decrease to 42 ± 4% of the pre-treatment value for the responsive hydrogel, while the inert hydrogel remained at 89 ± 2% (Figure 2B). After hydrogel deswelling, we investigated hydrolysis by trypsin. While inert hydrogels were visibly unaffected, responsive hydrogels disintegrated significantly (Figure S2C). As shown in Figure 2C, weight analysis revealed a post-treatment mass of 107 ± 9% and 15 ± 5% for inert and disintegrated responsive gels, respectively (for absolute values see **Table S2**).

To analyze whether deswelling and disintegration was caused by changes in the mechanical properties of the hydrogel, we tracked the storage modulus G’ by small amplitude oscillatory shear (SAOS) measurements.^[15,64]^ The parameters chosen for rheological characterization were validated to be within the linear viscoelastic range (LVE) via amplitude and frequency sweeps (**Figure S3**). As depicted in Figure 2D, enzymatic crosslinking by tyrosinase increased the storage modulus G’ of the responsive gels to 238 ± 38% (2.4-fold change), and of inert gels to 124 ± 8%, of their respective initial values. Subsequent trypsin hydrolysis disintegrated responsive hydrogels, preventing further analysis, while inert hydrogels reverted in storage modulus to 84 ± 12% of their initial state (not significant, p-value = 0.15). We hypothesize that the minor differences in weight and mechanical properties observed for inert hydrogels are caused by accumulation and subsequent removal of melanin-like polymers within the hydrogel matrix.

#### 2.2.2. Reversible shape-morphing by consecutive tyrosinase- and trypsin-mediated hydrogel modulation

To translate the enzyme-mediated volumetric changes of the hydrogels into controlled shape-morphing, we manufactured bilayer hydrogels by first forming an inert hydrogel layer which was subsequently overlayed with a responsive hydrogel. Here, the responsive layer was expected to undergo stronger deswelling than the inert one, in response to tyrosinase-mediated crosslinking. This process would force the bilayer material into a directional bending. Subsequent addition of trypsin as hydrolyzing enzyme would cleave hydrogel forming crosslinks in the responsive, but not the inert layer, thus reversing the bending motion through the elastic restoring force of the inert layer (**Figure 3A**).

**Figure 3.**
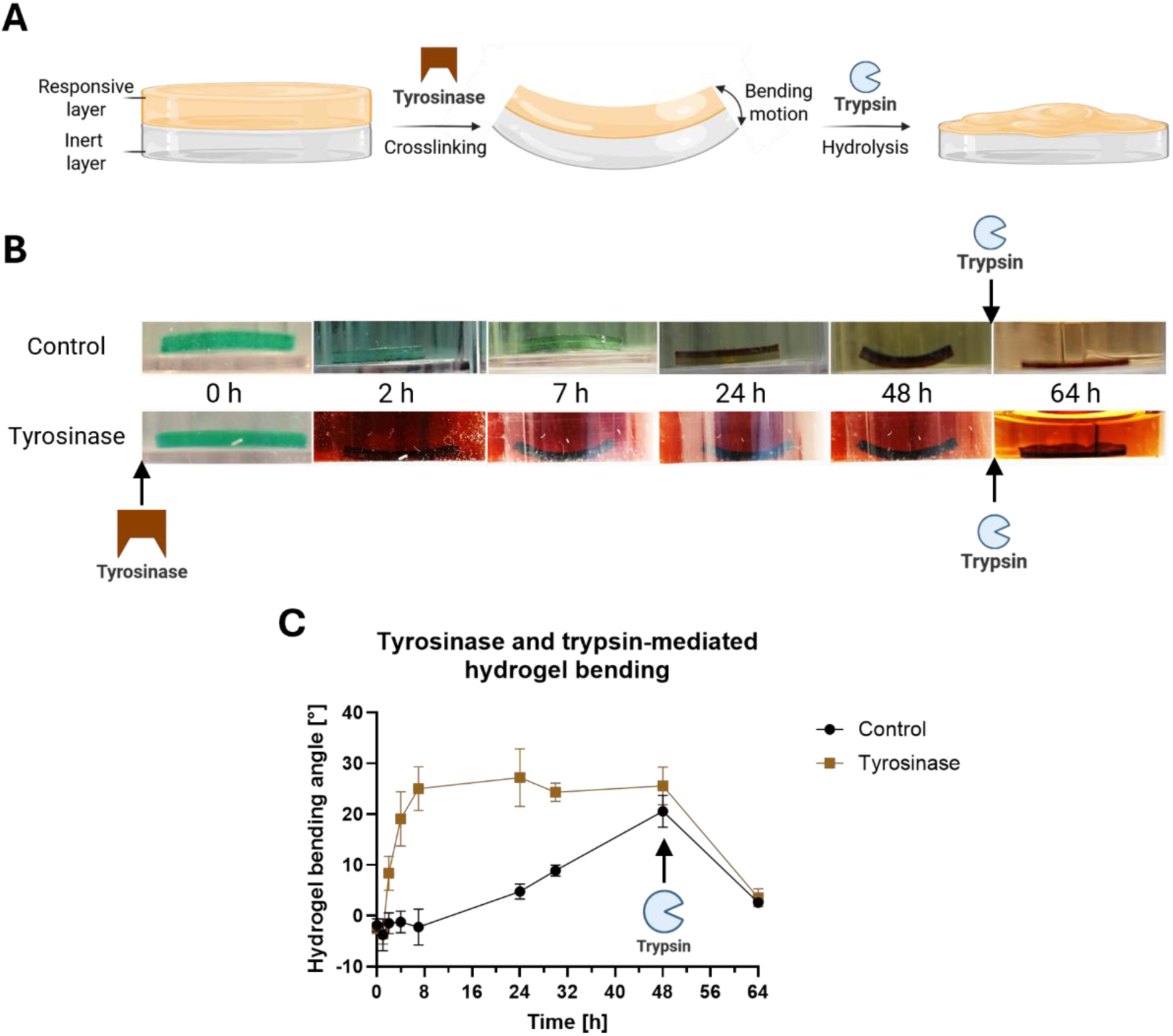
Tyrosinase- and trypsin-induced reversible shape-morphing of hydrogels. (A) Schematic representation of the bilayer hydrogel structure and bending mechanism. Upon crosslinking of the responsive layer by tyrosinase and CA, locally confined deswelling causes the bilayer hydrogel, consisting of an inert and responsive layer, to bend. Subsequent treatment with trypsin hydrolyzes bonds within the responsive layer, locally disintegrating the hydrogel and reversing the deformation. (B) Imaging of enzyme-mediated hydrogel bending. Bilayer hydrogels were generated by crafting the responsive layer on top of an inert hydrogel, each with a thickness of 0.5 mm and polymerization time of 20 min. Rectangular-shaped gels (2 x 10 mm) were cut and placed in M9 medium supplemented with green food dye for 2 h prior to the bending assay to enhance visibility. Subsequent enzyme treatments with tyrosinase or trypsin were performed as described in Figure 2B and 2C with control gels in M9 medium containing CA. Images were taken regularly to track the mechanical bending motion of the hydrogels. Depicted images are representative of replicates (n = 3) and were processed to enhance quality (sharpen, brightness and contrast); original images can be found in Figure S4. (C) Bending angle analysis of enzyme-mediated hydrogel deformation. Inner bending angles of the images from (B) were analyzed via the imaging software Fiji. Values are shown as mean ± SD (n = 3).

To test this concept, an inert hydrogel was prepared (0.5 mm height) and polymerized for 20 min prior to adding a second layer of equal height of the responsive hydrogel. The resulting temporary overlap of the polymerization processes in both layers enabled a covalent attachment of both due to the shared polymerization chemistry. Subsequently, rectangular shapes of 2 x 10 mm dimension were cut out and equilibrated in M9 medium overnight. The next day, images of the bilayer gels were taken, revealing an initial bending angle of −2 ± 1°. Here, the negative sign indicates a bending toward the inert layer, caused by the locally confined deswelling of the inert gel in this buffer (Figure S2A). Next, the hydrogels were incubated for 48 h in the presence of tyrosinase and CA and images were taken at indicated points in time (Figure 3B). We found that 7 h post treatment the responsive and control gels showed a significant bending angle difference of 27 ± 6° due to the enzyme-mediated increased bending rate. However, after 48 h untreated hydrogels had reached similar levels in bending and reduced the difference to 5 ± 5° (Figure 3C). We hypothesized that enzyme-independent CA autoxidation (indicated by hydrogel coloration), was responsible for hydrogel crosslinking at a slower rate, causing a delayed bending motion.^[65,66]^ Upon trypsin treatment, disintegration of the responsive layer caused bending angles to significantly decrease in tyrosinase-treated and non-treated hydrogels to 3 ± 1° and 4 ± 2°, respectively (Figure 3B and 3C). Together, these results demonstrate that laminating hydrogel layers with different dimensional changes upon enzymatic cues translates the underlying molecular mechanism into a directional and reversible bending motion of the bilayer material.

#### 2.2.3. Genetically programmed shape-morphing of bacteria-based ELMs

Next, we set out to design a bacteria-based ELM with genetically programmed shape-morphing. To this aim, we engineered *E. coli* to produce tyrosinase and incorporated them into the responsive layer to induce crosslinking, deswelling, and shape-morphing whereas subsequent trypsin addition would revert these processes (**Figure 4A**).

**Figure 4.**
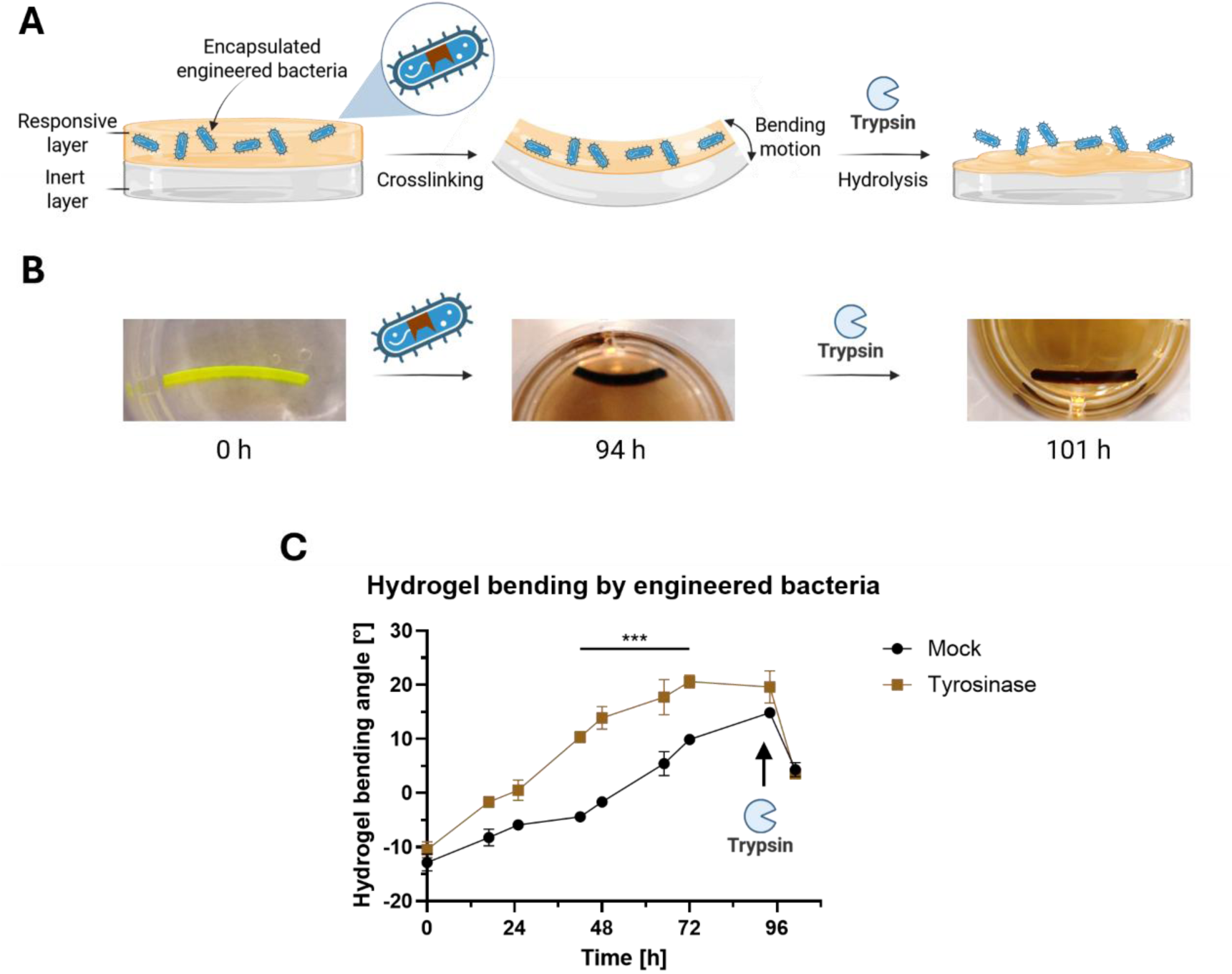
Bacterial ELMs with genetically programmed reversible shape-morphing. (A) Schematic depiction of hydrogel bending mediated by engineered bacteria. *E. coli* genetically programmed to produce tyrosinase, are embedded within the responsive layer to initiate local deswelling via crosslinking and as a result bending of the material. Subsequent treatment with trypsin hydrolyzes bonds exclusively within the responsive layer, reversing the deformation. (B) Imaging of bacteria-mediated hydrogel bending. *E. coli* transformed with the tyrosinase- or control protein-encoding plasmid were grown overnight until an OD_600_ of 0.8 – 1 was reached. To construct bilayer hydrogels, the inert layer of the hydrogel was prepared as described previously. Then, the responsive layer, modified by embedding of engineered bacteria (OD_600_ = 0.6) and integration of a fluorescent peptide to facilitate visual tracking, was added. After polymerization, rectangular-shaped (2 x 10 mm) hydrogels were equilibrated at 37 °C and 100 rpm for 2 h in M9, at which time-point CA and isopropyl-*β*-D-thiogalactopyranoside (IPTG) were added to induce tyrosinase production. Incubation of living hydrogels continued for 94 h with regular imaging to track mechanical deformation of the hydrogels. Afterward, hydrogels were incubated in trypsin (5000 U L^-1^) for 16 h and imaged. Depicted images are representative for replicates (n = 3) and were processed to enhance quality (sharpen, brightness, contrast); original images can be found in Figure S5C. (C) Bending angle analysis of bacteria-mediated hydrogel deformation. Inner bending angles of the images were analyzed via the imaging software Fiji. Values are shown as mean ± SD (n = 3), statistical significance was assessed by two-way ANOVA; ns = not significant, * = p ≤ 0.033, ** = p ≤ 0.002, *** = p ≤ 0.001.

We hypothesized that extracellular CA-mediated crosslinking should be possible despite the intracellular localization of the enzyme due to the inferred membrane permeability of CA.^[67–70]^ To test this hypothesis, we first analyzed extracellular CA conversion rate by intracellularly produced tyrosinase via Liquid Chromatography coupled to Electrospray Ionization Quadrupole Time-Of-Flight Mass Spectrometry (LC-ESI-QTOF-MS). Time-dependent substrate concentration differences were determined to calculate an intracellular enzymatic activity of 32.7 ± 3.4 U L^-1^ (Equation 1), for a detailed characterization including autoxidation rate see **Figure S5A**. To manufacture the ELM, *E. coli* transformed with a tyrosinase expression unit were integrated into a responsive hydrogel during polymerization. As shown in Figure S5B, the combination of CA and tyrosinase producing bacteria decreased hydrogel volume significantly to 58 ± 3% of the initial value, exceeding the deswelling caused by individual components. To translate the volumetric changes into shape-morphing, we prepared bilayer hydrogels as described above and restricted encapsulation of the tyrosinase-producing bacteria to the responsive layer. Further, for better identification of the layers, we applied the same PEG/peptide reaction chemistry to covalently attach a fluorescent (5,6-FAM) peptide into the responsive layer. As a control, *E. coli* transformed with an expression unit for an enzymatically inactive protein (DNA gyrase subunit B, GyrB) was used.^[71]^ The selected protein is functionless with respect to the performed assays but introduces a comparable metabolic burden. Again, CA was added as substrate to the ELMs, and hydrogel deformation was followed for 94 h (Figure 4B). Analysis of the bending angles revealed a significant difference between control-and tyrosinase-transformed bacteria during the time span of 36 to 72 h, with a maximum difference of 16 ± 2° after 48 h demonstrating the enzyme-mediated increased bending rate. (Figure 4C). As observed previously, prolonged incubation time (94 h) reduced the hydrogel deformation difference to 5 ± 3. We hypothesize that the substantial bending of the mock group was driven by CA autoxidation, indicated by the brown coloration of the ELM with the control protein (Figure S5C). Subsequent trypsin treatment reversed the bending angles to 4 ± 2° and 4 ± 1° for GyrB and tyrosinase expressing hydrogels, respectively (Figure 4B and 4C). Taken together these data demonstrate that the ELM can be genetically programmed to autonomously perform defined shape-morphing. Further, the ELM features an external, trypsin-mediated control mechanism allowing the overriding or the reversion of the genetically induced mechanical action.

### 2.3. Transglutaminase and PPEP-1-mediated hydrogel modulation

To expand the actuator toolbox and demonstrate the flexibility of our approach for different applications, we next combined the crosslinking enzyme microbial transglutaminase (TG) with the metalloprotease PPEP-1 for a more selective bond formation and hydrolysis.

#### 2.3.1. Transglutaminase and PPEP-1-mediated reversible swelling and mechanical properties changes

Since TG introduces crosslinks between K and glutamine (Q) residues (**Figure 5A** and S1D), we formed the hydrogel by crosslinking 8-arm PEG-VS with an equimolar mix of two peptides: Peptide-K (see above) and a peptide containing a single Q residue (Peptide-Q). Cleavage of the peptide can be performed with high specificity by the metalloprotease PPEP-1 hydrolyzing a specific proline-proline (P-P) bond within the peptide sequence PNP↓PVP (Figure 5A and S1E).^[72]^ This process leaves the isopeptide bond introduced by TG intact to stabilize the hydrogel network, resulting in a partial network hydrolysis and therefore swollen, but not disintegrated, hydrogel. As before, the inert hydrogel crosslinked by DTT was used as control. Like above, responsive and inert hydrogels were formed and equilibrated in phosphate buffered saline (PBS, pH 7.4) showing an equilibration behavior to 99 ± 2% and 144 ± 4% of polymerized weight, respectively (**Figure S6A**). After subjecting the hydrogels to TG (1400 U L^-1^) treatment, the hydrogel size of the responsive gel decreased visibly, while the inert sample remained unchanged (Figure S6B). Analysis of the hydrogel masses revealed a decrease to 59 ± 5% of pre-treatment weight for responsive gels, and 98 ± 2% for inert hydrogels (Figure 5B). The crosslinked hydrogels were subsequently incubated with PPEP-1 (0.48 U L^-1^, **Figure S7**), revealing a significant swelling (Figure S6C) to 183 ± 24% for responsive gels while inert gels remained at 92 ± 6% (Figure 5C, for absolute values see **Table S3**). As described above, SAOS measurements were performed within the LVE range (**Figure S8**). G’ determination revealed that the storage modulus of the responsive hydrogel increased to 554 ± 140% (5.5-fold change) upon TG treatment, while the inert version increased less significantly to 147 ± 8%. We attribute the latter observed effect to self-crosslinking of TG via intrinsic K and Q residues, causing the enzyme to entangle within the hydrogel matrix and therefore altering the physical properties of the polymer network.^[73,74]^ After subsequent protease treatment, the responsive gel decreased to 206 ± 65% of the initial G’ while the inert gel remained at 147 ± 20% (Figure 5D).

**Figure 5.**
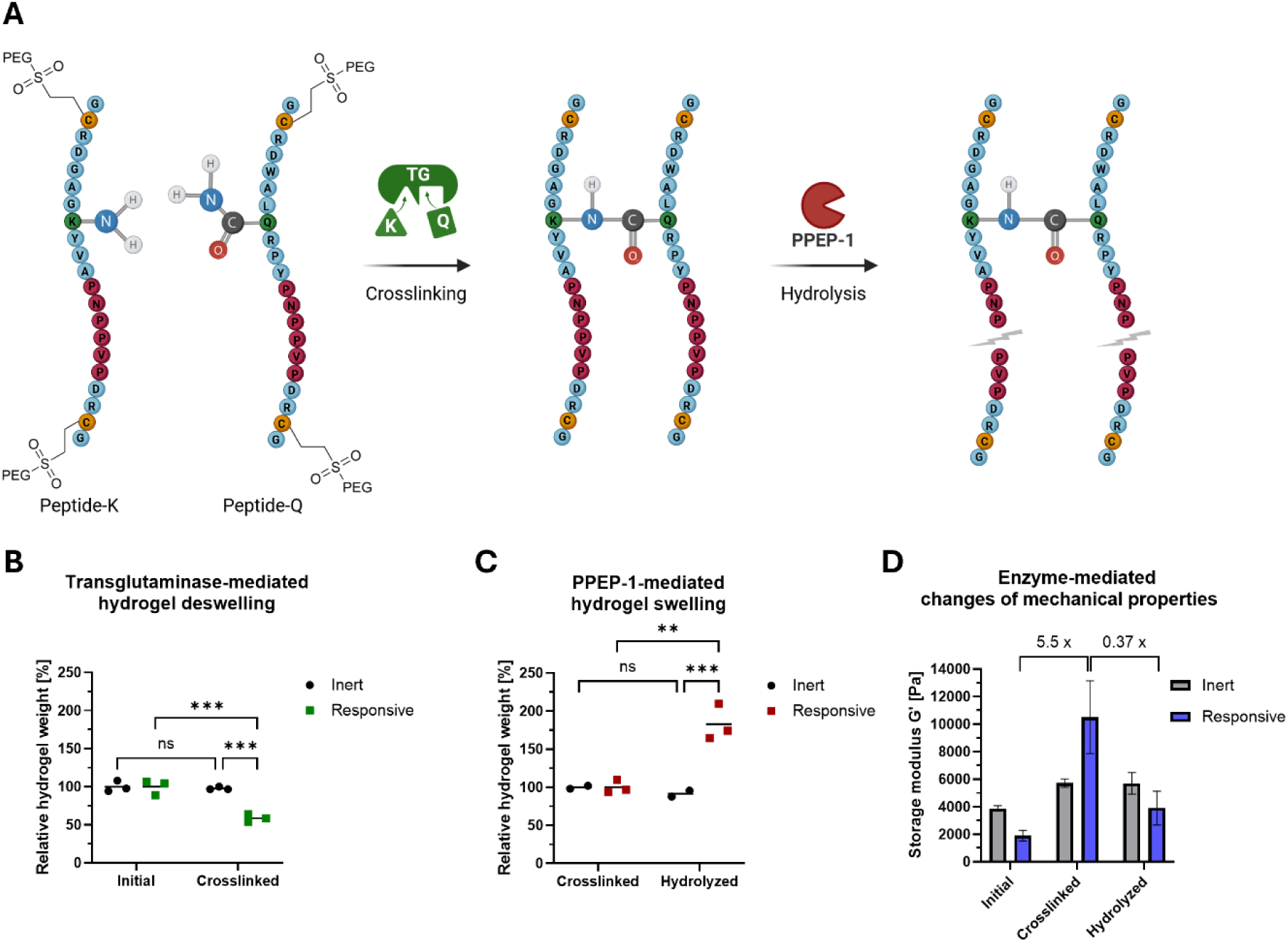
Mechanical and physical characterization of transglutaminase- and PPEP-1-responsive hydrogels. (A) Schematic representation of enzyme-mediated network modulation. Transglutaminase (TG)-mediated crosslinking by isopeptide bond formation between K and glutamine (Q) residues contained in the peptides (Peptide-K and Peptide-Q) used to form a hydrogel with 8-arm PEG-VS. Upon addition of the metalloprotease PPEP-1, proline-proline (P) peptide bonds are hydrolyzed, reducing crosslinking density and inducing hydrogel swelling. (B) Transglutaminase-mediated hydrogel deswelling. Hydrogels were prepared as depicted in Figure 1B: 8-arm PEG-VS was crosslinked with either DTT to form biologically inert hydrogels, or with a 1:1 molar ratio of K/Q containing peptides to form biologically responsive hydrogels. After 60 min polymerization, hydrogels were equilibrated in phosphate buffered saline (PBS) overnight and subsequently initial weight was recorded. Afterwards, crosslinking was initiated by TG addition (1400 U L^-1^) and continued for 24 h at 37 °C before hydrogel mass was recorded. (C) PPEP-1-mediated hydrogel swelling. Crosslinked gels from (B) were washed and subsequently incubated with PPEP-1 (0.48 U L^-1^) at 37 °C and further mass changes recorded after 16 h. (D) Enzyme-mediated changes in mechanical properties of hydrogels. G’ of hydrogels from (B) and (C) were determined via rheological measurements before and after enzymatic treatments. Values are shown as mean ± SD (n = 2 - 3). Normalization for hydrogel weight was performed relative to pre-treatment value, statistical significance was assessed by two-way ANOVA; ns = not significant, * = p ≤ 0.033, ** = p ≤ 0.002, *** = p ≤ 0.001. One sample was inadvertently damaged during processing and therefore excluded from the analysis and corresponding Figure (C) and (D).

#### 2.3.2. Reversible shape-morphing by consecutive transglutaminase- and PPEP-1-mediated hydrogel modulation

Next, we set out to analyze the reversible shape-morphing potential mediated by the TG and PPEP-1 pair. Upon treatment of bilayer hydrogels with TG, crosslinking confined to the responsive layer was expected to initiate a bending motion towards this side. Conversely, subsequent PPEP-1 hydrolysis would reduce network density in the responsive layer, reversing hydrogel actuation in a controlled manner (**Figure 6A**). Bilayer hydrogels with an equimolar peptide mixture (Q, K) in the responsive layer were fabricated and equilibrated in PBS. Untreated gels showed an initial bending angle of 26 ± 2° toward the responsive layer, in line with our previous results (Figure S6A) showing an increased swelling for the inert hydrogels in this buffer. Afterwards, gels were treated with TG to induce crosslinking of the responsive layer, while control bilayer gels were kept in PBS. As shown in Figure 6B and 6C (left part), enzyme-treated bilayer gels significantly deformed in a bending motion toward the responsive layer, with a final bending angle of 112 ± 6°, while untreated samples showed no changes (26 ± 4°). To evaluate reversible bending, crosslinked hydrogels were incubated in the presence or absence of PPEP-1 (0.48 U L^-1^). While control samples only revealed a minor decrease in bending angle from 106 ± 3 to 88 ± 7°, PPEP-1-treated bilayer gels showed a significant decrease from 116 ± 1 to 45 ± 3° (Figure 6B and 6C, right part). We anticipate that constant strain exerted by the inert layer, passive hydrolysis of peptide bonds, as well as removal of TG from the responsive matrix are responsible for a mild relaxation of the control bilayer hydrogel. This reversible actuation could also be applied to a more intricate shape such as a cross-shaped hydrogel. Sequential exposure of this complex shape to TG and PPEP-1 demonstrated a strong shape-morphing response imitating reversible closing and opening of a blossom (Figure 6D, Supplementary Movie 1).

**Figure 6.**
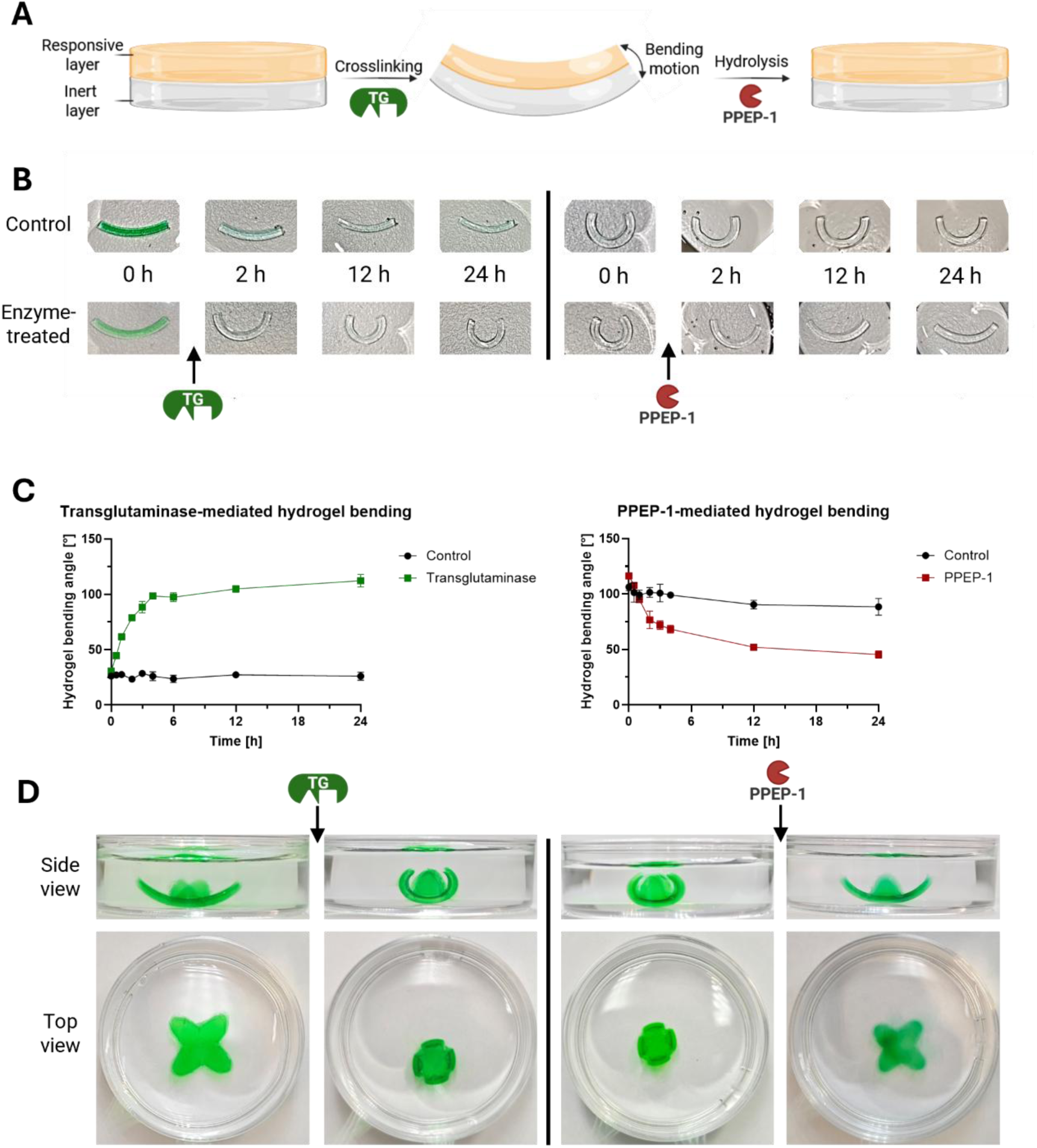
Transglutaminase- and PPEP-1-induced reversible shape-morphing of hydrogels. (A) Schematic representation of the bilayer hydrogel structure and bending mechanism. Upon crosslinking of the responsive layer by TG, locally confined deswelling causes the material to bend. Subsequent treatment with the protease PPEP-1 hydrolyzes crosslinks within the responsive layer, reversing the deformation. (B) Imaging of enzyme-mediated hydrogel bending. Bilayer hydrogels were generated as described previously. After polymerization, hydrogels were equilibrated in PBS supplemented with green food dye to enhance visibility. Subsequent enzyme treatments with TG and PPEP-1 were performed sequentially as described before at the indicated timepoints with untreated gels as controls. Images were taken regularly to track the mechanical bending motion of the hydrogels (B). Depicted images are representative for replicates (n = 3) and were processed to enhance quality; for original images see Figure S9. (C) Bending angle analysis of enzyme-mediated hydrogel deformation. Inner bending angles after TG (left) or PPEP-1 (right) treatment of hydrogels from the images in (B) were analyzed via the imaging software Fiji. Values are shown as mean ± SD (n = 3). (D) Enzyme-mediated complex shape-morphing. Cross-shaped bilayer hydrogels were fabricated by punching with a suitable shape (peak-to-peak distance 1.3 mm) and first treated with TG (1400 U L^-1^) for 6 h before being submitted to PPEP-1 hydrolysis (2.88 U L^-1^) for 6 h.

#### 2.3.3 Genetically programmed out-of-equilibrium ELM with autonomous, reversible shape-morphing

Out-of-equilibrium materials are materials in which counteracting chemical reactions are constantly modifying the material’s properties and where the material’s steady state depends on the relative rates of these counteracting reactions. Such materials have shown fast and dynamic responsiveness to external or material-internal stimuli modulating the relative reaction rates.^[75,76]^ Here, we apply the above-established TG- and PPEP-1-mediated processes to devise an out-of-equilibrium ELM, the shape-morphing of which was controlled by opposing biochemical reactions with a genetically programmed dynamically changing reaction rate. To this aim, we engineered mammalian cells to produce and secrete PPEP-1 and placed the cells onto TG/PPEP-1-responsive bilayer gels (**Figure 7A**). We further added TG from the beginning. In this configuration, we expect simultaneous TG- and PPEP-1-mediated crosslink formation and hydrolysis of the responsive layer. PPEP-1-mediated hydrolysis would be expected to increase over time due to accumulation of the protease while TG-induced crosslinking would remain constant, decrease over time or face substrate-depletion. This genetically encoded shift in relative reaction rates is expected to result in autonomous reversible actuation with an initial, TG-dominated bending and a subsequent, PPEP-1-dominated relaxation (Figure 7A).

**Figure 7.**
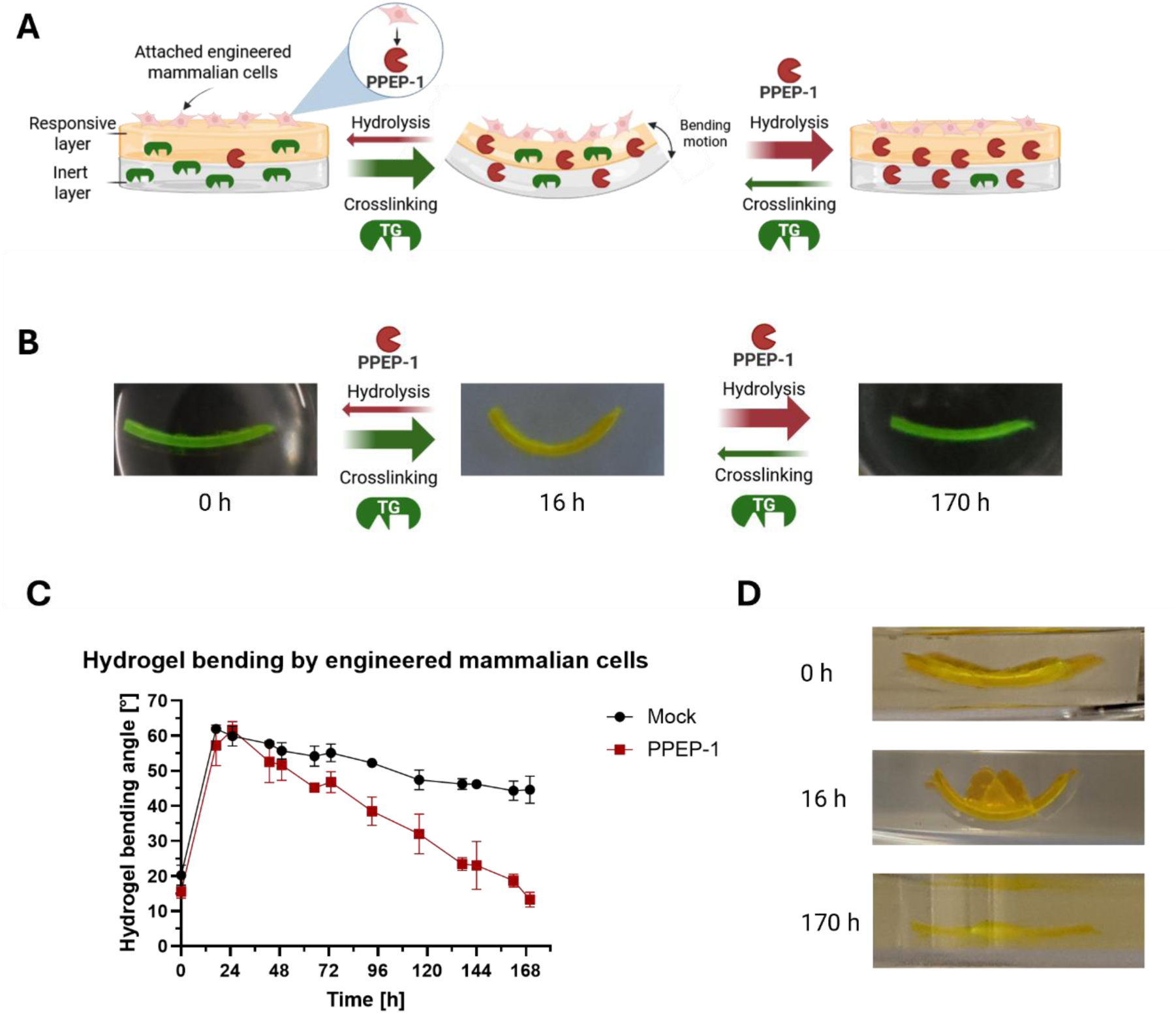
Autonomous reversible shape-morphing of a mammalian ELM controlled by out-of-equilibrium counteracting biochemical reactions with dynamically changing relative reaction rates. (A) Schematic depiction of hydrogel bending by mammalian cells. TG and HEK-293T engineered to secrete PPEP-1 are added simultaneously to a bilayer hydrogel. A fluorescent RGD motif containing peptide within the responsive layer, allows for incorporation of mammalian cells onto the hydrogel and enhanced visual tracking. Initially, TG activity dominates, leading to crosslinking of the responsive layer and initiates a bending of the hydrogel. Over time, the cell-secreted enzyme PPEP-1 accumulates and reverses the deformation by decreasing crosslinking density in the responsive hydrogel layer. (B) Imaging of cell-mediated hydrogel deformation. Bilayer hydrogels were prepared as described previously and equilibrated for 2 h in complete cell culture medium prior to cell seeding. HEK-293T engineered to produce PPEP-1 or control cells containing an empty vector were seeded to the hydrogels and TG (1400 U L^-1^) was added. Hydrogels were incubated for 170 h and images were taken regularly to track hydrogel bending. Depicted images are representative for replicates (n = 3) and were sharpened to enhance quality; original images can be found in Figure S10. (C) Bending angle analysis of cell-mediated hydrogel deformation. Inner bending angles of images were analyzed via the imaging software Fiji. Values are shown as mean ± SD (n = 3). (D) Enzyme-mediated complex shape-morphing. Cross-shaped bilayer hydrogels (peak-to-peak distance 0.9 mm) were prepared and treated as described in (B). Images were taken at the indicated time-points to track hydrogel bending.

To enable PPEP-1 secretion, the enzyme-encoding sequence was fused to the human interleukin-6 (IL-6)-derived secretion sequence. The resulting construct was placed under the control of the strong CMV promoter and transfected into HEK-293T cells, revealing a PPEP-1 production rate of 0.20 ± 0.09 U (h * 10^6^ cells)^-1^ (Equation 2 and 3). Second, we modified the hydrogel to support adhesion of mammalian cells and enable visual tracking. To this aim, a fluorescent peptide harboring the RGD motif to interact with the cells’ integrins was incorporated into the responsive hydrogel layer.

We used this system to analyze autonomous shape-morphing controlled by counteracting and dynamically changing TG and PPEP-1 reaction rates. To this aim, bilayer gels were synthesized, seeded with PPEP-1 producing HEK-293Tcells (2 x 10^5^ cells per gel) and supplemented by TG (1400 U L^-1^) prior to following the hydrogel deformation over 170 h. As shown in Figure 7B, the bilayer gels first showed strong bending followed by a relaxation phase. Quantitative analysis revealed that hydrogels with mock or PPEP-1 transfected cells showed a respective bending angle increase of 42° ± 2 or 42° ± 4 after 16 h. Subsequently, the relaxation phase started, characterized by a reversion of the bending by 44 ± 6° for the PPEP-1 producing ELMs compared to only 17 ± 4° for mock-transfected cells (Figure 7C). In addition to the previously observed mild relaxation of control gels (Figure 6C), this effect could be attributed to the production of non-specific hydrolytic enzymes of the cells capable of cleaving the peptide, which was determined to be 0.28 ± 0.10 U (h * 10^6^ cells)^-1^. We further demonstrated that this autonomously timed reversible shape-morphing could be used to control more complex shape trajectories such as a cross-shaped bilayer hydrogel (Figure 7D). These data demonstrate the potential of using genetically programmed biochemical reaction rates to control autonomous macroscopic shape-morphing of ELMs.

## 3. Discussion

In this work, we develop ELMs with genetically and enzymatically programmed autonomous and reversible bending. The ELMs are based on a biologically responsive hydrogel whose volumetric dimensions and mechanical properties can be dynamically and reversibly modified via two antithetic enzyme pairs with distinct specificities and mechanisms. By fabricating bilayer hydrogels with asymmetric responsivity, these molecular transformations were translated into a directed macroscopic motion. The integration of genetically programmed or external supplementation with enzymes was shown to program autonomous and reversible motion with counteracting reactions.

The presented synergy of genetic programming and material design offers a versatile strategy for autonomous ELM actuation, addressing current limitations of systems reliant on externally applied stimuli such as heat, light, or pH.^[26,27,48]^ By translating genetic information into mechanical output, material shape-morphing becomes pre-programmable and executable, advancing stimulus-responsive materials towards genuine self-regulation and autonomy. Further, the dual responsiveness of the presented ELMs enables the generation of out-of-equilibrium ELMs, that allow fast and dynamic responses and programming by tuning the relative reaction rates of the opposing enzymatic reactions.

Conceptually, this approach complements existing cell-based actuation platforms in which motion arises from cellular contraction or proliferation.^[32,49,50,77]^ While mammalian cell guided contraction only operates in a narrow window of environmental conditions, thereby limiting application potential, a controlled reversibility of proliferation-controlled systems remains challenging. Recent advances have addressed the latter by externally adding enzymes to degrade yeast cell walls, achieving partial shape-recovery and enabling subsequent growth/degradation cycles.^[78]^ Mechanistically, such approaches distinguish themselves from this work since they employ material content modification to achieve volumetric changes, while enzymatic shape-morphing is mediated through network remodeling. These cell-produced actuators operate as self-contained molecular machines with built-in programmability across several stages of actuation (e.g., expression, production, activity, substrate availability), further allowing combinatorial application of actuators to achieve dynamic steady state or targeted sequential modulations.

While the enzyme-mediated bending motions demonstrated in this study deliver meaningful mechanical outputs and fall within ranges of soft robotic bending elements, several challenges remain to harness the full potential of this concept.^[47,48]^ The time frame and extent of material deformation depend on the lifetime and activity of the integrated cells and produced actuators, as well as enzyme reaction kinetics and diffusion limitations within the material. Although the applied opposing enzymes can reverse the macroscopic deformation through orthogonal mechanisms, the molecular architecture is not returned to its original state, causing substrate depletion to limit motion cycling. Furthermore, despite the reproducibility of PEG–peptide hydrogel manufacturing, scalable fabrication workflows of complex, cell embedding shapes remains a considerable engineering challenge.^[5]^

Despite these limitations, the translation of genetically programmed information into a reversible shape-morphing within a fully synthetic hydrogel marks a substantial step towards autonomous soft robotics, self-regulating biohybrid actuators, and dynamic microfluidic systems. Going forward, wiring actuator production to molecular biosensors and gene circuits will enable ELMs to sense, process, and respond mechanically to environmental cues. Such integration not only reduces reliance on external hardware but also facilitates genetically programmable logic and feedback like threshold responses, adaptation, and oscillations, paving the way for the next generation of out-of-equilibrium shape-morphing ELMs. These self-regulatory systems could ultimately be employed to detect biochemical signals and serve as drug depots that mechanically deliver therapeutics, adaptive biomedical implants that modulate their stiffness and geometry, and environmental pathogen sensors that report status changes through mechanical deformation.

## 4. Experimental Section

### Hydrogel synthesis

8-arm 10 kDa MW PEG-vinyl sulfone (PEG-VS, SKU: A10033-1) was purchased from JenKem Technology USA. Custom fluorescent RGD motif containing peptide ((5,6-FAM)-GCGYGRGDSPG-NH_2_) and crosslinking peptides (Ac-GCRDWALQRPYPNPPVPDRCG-NH_2_ (Peptide-Q) and Ac-GCRDGAGKYVAPNPPVPDRCG-NH_2_ (Peptide-K)) were synthesized by Caslo ApS (Technical University of Denmark). Peptide sequences are written N-terminus to C-terminus with acetylated (Ac) and amidated (NH_2_) termini. All reagents were dissolved in 0.3 M triethanolamine (TEA, pH 8, Carl Roth, art. no. 6300.1) and used directly or stored at −80 °C until required. A detailed overview of stock concentrations and hydrogel compositions can be found in **Table 1**. For hydrogel preparation, PEG-VS was first diluted with TEA and optionally RGD peptide was added. The solution was vortexed briefly, and if RGD was present, allowed to react for 5 min at room temperature (RT). Afterwards, crosslinking peptides (Peptide-K and Peptide-Q) or DTT were added in a 1.25 molar excess of thiol to available VS groups. For gels for subsequent tyrosinase treatment, only Peptide-K was used, while hydrogels used for TG crosslinking contained an equimolar mixture of Peptide-K and Peptide-Q. The hydrogel precursor was vortexed briefly again and transferred onto a hydrophobically coated glass slide (Sigmacote). Generation of disc-shaped gels was achieved by placing two spacers with 0.8 mm height adjacent to the gels and placing a second hydrophobic glass slide on top. The hydrogels were then polymerized for 60 min at RT. For bilayer hydrogels, individual layer thickness and polymerization time were reduced to 0.5 mm and 20 min, respectively. Rectangular (2 x 10 mm) or cross-shaped forms (peak-to-peak 0.9 or 1.3 mm as indicated) were used to cut out hydrogels for bending experiments.

**Table 1.**
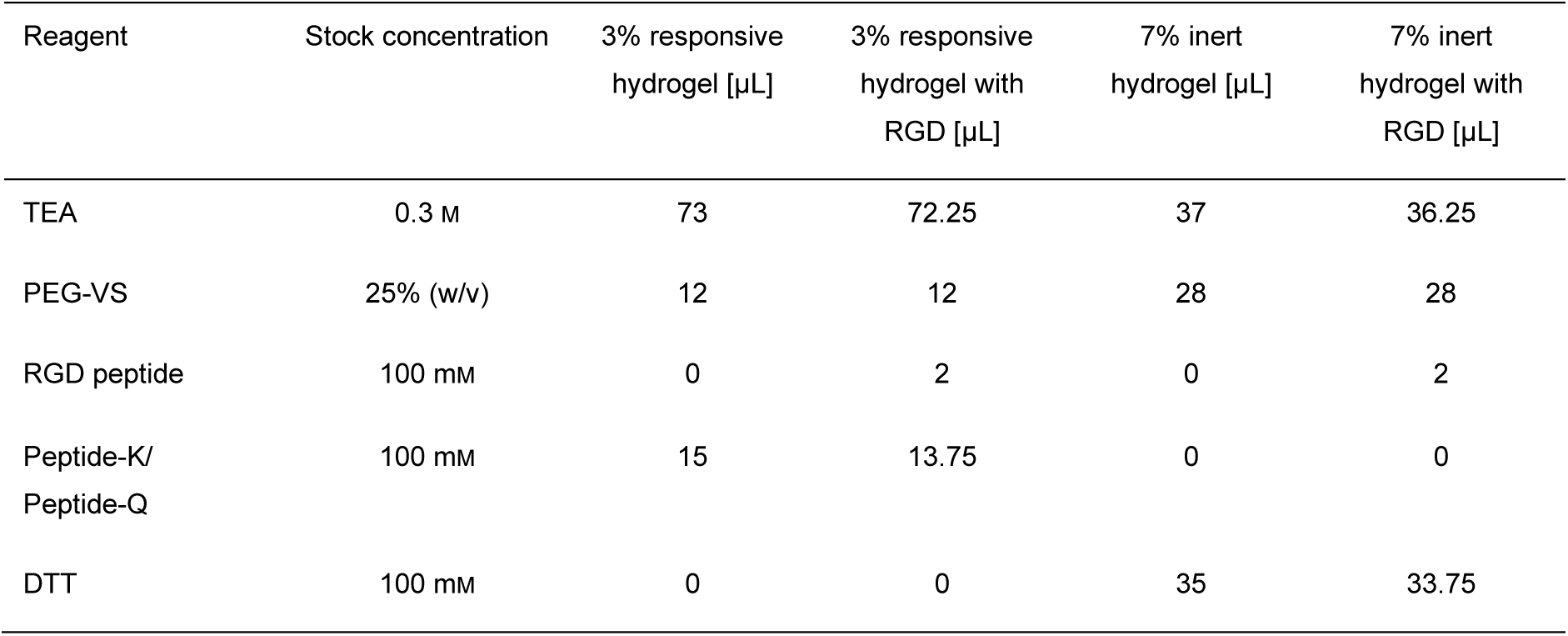
Hydrogel compositions for 100 µL final volume.

| Reagent | Stock concentration | 3% responsive hydrogel [μL] | 3% responsive hydrogel with RGD [μL] | 7% inert hydrogel [μL] | 7% inert hydrogel with RGD [μL] |
| --- | --- | --- | --- | --- | --- |
| TEA | 0.3 M | 73 | 72.25 | 37 | 36.25 |
| PEG-VS | 25% (w/v) | 12 | 12 | 28 | 28 |
| RGD peptide | 100 mM | 0 | 2 | 0 | 2 |
| Peptide-K/<br>Peptide-Q | 100 mM | 15 | 13.75 | 0 | 0 |
| DTT | 100 mM | 0 | 0 | 35 | 33.75 |

### Initial hydrogel swelling

After polymerization, tyrosinase-responsive hydrogels were washed with 2 mL minimal medium (M9; 48 mM NaH_2_PO_4_, 22 mM KH_2_PO_4_, 8.6 mM NaCl, 10 mM glucose, 1 mM MgSO_4_, 10 mM L-aspartate, 10 µM CuSO_4_, 1 mM ampicillin and chloramphenicol, pH 6.8) supplemented with L-cysteine (C, 10 mM) for 2 h. Afterwards, gels were equilibrated overnight in a C-free formulation of this medium. For TG crosslinking, the hydrogels were instead directly placed in phosphate buffered saline (PBS) for overnight swelling. Green food dye (Lianyungang Xinai Food Technoloy) was used in a 1:1000 dilution in the respective buffer to dye the hydrogels for visibility.

### Cloning of expression plasmids

All expression plasmids were prepared by Gibson assembly or restriction enzyme cloning. The respective DNA sequences are depicted in **Table S1** and were confirmed by Sanger sequencing. The tyrosinase sequence was taken from *Priestia megaterium*. For PPEP-1, the signaling peptide (AA 2-26) was removed from the original sequence derived from *Clostridium difficile*. For transfection of mammalian cells, the mammalian secretion signal derived from human IL-6 was inserted instead.

### Protein production and purification

Tyrosinase was produced in *E. coli* (BL21 DE3 pLysS, Invitrogen) transformed with plasmid pRS39 in LB medium supplemented with ampicillin (100 µg mL^-1^). Following induction with 1 mM isopropyl-*β*-D-thiogalactopyranoside (IPTG) at OD600 of 0.6 - 0.8, production was performed at 18 °C and 150 rpm overnight. Cells were harvested by centrifugation (6000 g, 10 min, 4 °C) and subsequently frozen in liquid nitrogen before storage at −80 °C. Cell pellets were thawed and resuspended as 20% (w/v) in Ni-Lysis buffer (50 mM NaH_2_PO_4_, 300 mM NaCl, 10 mM imidazole, pH 8.0) before incubation with lysozyme (∼1 mg, Carl Roth, art. no. 8259) for 1 h at 4 °C under agitation. Afterwards, cell lysis was performed with a high-pressure homogenizer (20 kpsi, Multi Cycle Cell Disruptor, Constant Systems Ltd.) and the resulting suspension was centrifuged (30000 g, 30 min, 4 °C). The cleared supernatant was loaded and purified via immobilized metal affinity chromatography (IMAC) as described previously.^[79]^ After IMAC purification, a buffer exchange to sodium phosphate buffer (50 mM, pH 6.7) was performed by repeated centrifugation (6000 g) and resuspension via a concentrator membrane (Vivaspin 10 kDa CO, ref. VS0601). Glycerol was added (15% (v/v) final concentration) and protein was stored at - 80 °C.

For PPEP-1 production, *E. coli* cells (BL21 DE3 pLysS) were freshly transformed with the plasmid pJB001 and cultivated overnight at 37 °C and 180 rpm in LB medium supplemented with ampicillin (100 µg mL^-1^). On the next day, the culture was inoculated and grown until an OD_600_ of 0.6 - 0.8 was reached prior to expression induction by IPTG (1 mM). Production was performed for 5 h and cells subsequently harvested by centrifugation (6500 g, 15 min, 4 °C). After elimination of the supernatant, bacteria from 1 L culture were resuspended with 40 mL Ni-Lysis buffer and shock-frozen in liquid nitrogen. For purification, the frozen cell suspension was thawed at 37 °C, sonicated (15 min, 40%) and centrifuged (30000 g, 30 min, 4 °C). After equilibration of a gravity flow column (ThermoScientific, ref. 29924), containing 1 column volume (CV, 1 CV = 2 mL) Ni-nitrilotriacetic acid beads (Qiagen, ref. 30430), with 5 CV of Ni-Lysis buffer, the cleared lysate was applied. The column was then washed with 10 CV Ni-Wash buffer (50 mM NaH_2_PO_4_, 300 mM NaCl, 20 mM imidazole, pH 8.0), followed by protein elution with 2 CV of Ni-Elution buffer (50 mM NaH_2_PO_4_, 300 mM NaCl, 250 mM imidazole, pH 8.0). Eluted protein was stored at −80 °C until further use.

### Protein characterization

PPEP-1 samples were incubated in 1x SDS loading buffer (from 5x stock solution with 50% (v/v) glycerol, 312.5 mM Tris, 12.5% (v/v) 2-mercaptoethanol (2-ME), 10% (w/v) SDS, 0.05% Bromophenol blue) at 95 °C for 5 min and subsequently loaded onto an 10% sodium dodecyl sulfate-polyacrylamide gel electrophoresis (SDS-PAGE) gel. Afterwards, proteins within the gel were stained with Coomassie blue solution, revealing a band at the expected size (24.2 kDa, Figure S7). Spectrophotometry measurement of PPEP-1 revealed a protein concentration of 1.27 mg mL^-1^. The SDS-PAGE analysis of tyrosinase revealed a distinct band at the expected size (35.2 kDa, data not shown). Final protein concentration was determined to be 1.5 mg mL^-1^ by microvolume spectrophotometry (Thermo Scientific, Nanodrop One^C^).

### Rheological characterization

For rheological measurements a parallel plate setup with sandblasted surfaces was used at a rheometer (Anton Paar MCR302e). To this aim, disc-shaped hydrogel diameter was equalized to a diameter of 8- or 12-mm with a biopsy punch. Afterwards, the gels were loaded onto a pre-heated (37 °C) lower plate (Anton Paar, P-PTD220/air/xx2) and 50 µl appropriate buffer was added to prevent desiccation. The sample was then compressed with the respective upper plate (PP08/S or PP12/S) to a constant normal force of 0.03 N and equilibrated for 2 min, before measurements were started. For amplitude sweeps, a constant frequency of 1 Hz and a ramp function for deformation (0.01 - 100%) was applied. Frequency sweeps were performed with a constant deformation of 0.1% and a frequency range (0.1 – 10 Hz). Finally, small amplitude oscillatory measurements (SAOS) were conducted with pre-determined values (0.1% and 1 Hz) within the LVE range.

### Hydrogel enzyme treatment

Tyrosinase-mediated crosslinking of the gels was performed in M9 with tyrosinase (0.05 mg mL^-1^) and caffeic acid (5 mM final concentration from 1.5 M stock in dimethyl sulfoxide) for 48 h at 37 °C with a medium exchange after 24 h. Crosslinking by TG was conducted for 24 h at 37 °C with an enzyme concentration of 10 mg mL^-1^ (1400 U L^-1^ (enzyme units per liter), Transglutaminase TI, 140 U g^-1^, Bindly) in PBS. After enzyme treatments, hydrogels were washed for 2 h in 2 mL of their respective buffer.

Tyrosinase-crosslinked gels were hydrolyzed with trypsin (Trypsin 0.05% (w/v), approximately 5000 U L^-1^, PAN Biotech Cat. No. P10-023500), while hydrogels crosslinked by TG were treated with PPEP-1 (0.1 mg mL^-1^), both at 37 °C for the indicated incubation period. For the accelerated PPEP-1 hydrolysis shown in Figure 5D and the time lapse video, a protease concentration of 0.6 mg mL^-1^ was used.

### Hydrogel weighing and imaging

After each step altering hydrogel dimensions (enzyme treatment or punching), hydrogel weight and/or diameter were documented. Hydrogel mass was determined with a precision scale (Mettler Toledo, XPR225) after carefully removing excess surface liquid. Images were taken with a digital camera (Canon, EOS R50) and inner bending angles of bilayer hydrogels were determined manually with the Fiji (Version 1.54p) angle tool.

### Bacteria transformation and cultivation

*E. coli* cells (BL21 DE3 pLysS) were transformed with the respective plasmids encoding GyrB or tyrosinase and cultivated for 1 h in 200 µL LB medium at 37 °C before being transferred into 10 mL M9 for overnight incubation (37 °C, 180 rpm). On the next day, bacteria were harvested by centrifugation (10 min, 6000 rpm, 4 °C) and the cell pellet was resuspended in an appropriate volume of M9 to reach a final OD_600_ of 0.6 - 0.8 as indicated in the respective experiments.

### Bacteria encapsulation and hydrogel bending

For bacterial bilayer hydrogel fabrication, the responsive layer was modified to contain the fluorescent RGD peptide (Table 1), and the TEA portion was replaced with bacterial cell suspension (OD_600_ = 0.8). The rectangular or cross-shaped bilayer gels were cut out and incubated for 1 h at 37 °C under shaking conditions, first in M9 supplemented with C (10 mM) and then in C-free M9. Afterwards, the solution was replaced with M9 (1.5 mL for rectangular, 3 mL for cross-shaped) containing CA (5 mM) and IPTG (1 mM) and incubation was continued as indicated in the respective experiments. To prevent desiccation, fresh M9 (1/3 of initial volume) was added after 48 h.

### Caffeic acid conversion rate determination by Liquid Chromatography coupled to Electrospray Ionization Time-Of-Flight Mass Spectrometry (LC-ESI-QTOF-MS)

After resuspension of bacteria to OD_600_ = 0.6, IPTG (1 mM) was added and bacteria grown for 5 h at 37 °C and 180 rpm. Afterward, cells were harvested by centrifugation (6000 rpm, 10 min, 4 °C) and resuspended in M9 to OD_600_ = 2. The concentrated cell suspensions were diluted in fresh M9 to reach a final OD_600_ of 0.6 in 500 µL and were then equilibrated for 10 min at 37 °C and 100 rpm, before CA (5 mM) was added. As references, 5 µg mL^-1^ purified tyrosinase and CA without enzyme were incubated. After 60 and 90 min incubation periods, samples were collected on ice and immediately centrifuged (8000 g, 15 min, 4 °C) to stop enzymatic activity. Supernatant was collected and stored at −20 °C until further analysis. For analysis of CA autoxidation, the incubation periods were adjusted to 60 min and 25 h.

LC-ESI-QTOF-MS analysis was performed on a 1260 Infinity LC in combination with a 6545A high-resolution time-of-flight mass spectrometer, both from Agilent Technologies (Santa Barbara, CA, USA). The separation of 0.5 µL samples was performed using a Poroshell HPH-C18 column (3.0 x 50 mm, 2.7 µm) equipped with the same guard column (3.0 x 5 mm, 2.7 µm) under isocratic conditions with 35% methanol in water containing 0.1% formic acid at a flow rate of 200 µL min^-1^ at RT.

After separation, the LC flow entered the dual AJS ESI source, which was set to 3000 V as capillary voltage and 500 V as nozzle voltage, 50 psi nebulizer gas pressure, 12 L min^-1^ dry gas flow, and 350 °C dry gas temperature. The MS parameters used were 2 GHz (extended dynamic range), 150 V fragmentor voltage, and 45 V skimmer voltage. The mass spectra were recorded between 0.5 - 4 min in full scan mode in the mass-to-charge ratio (*m/z*) range of 100 - 1000 Da at a spectrum rate of 1 s^-1^.

For quantification, the negatively charged exact mass [M-H] for CA (C_9_H_8_O_4_) at *m/z* 179.036 Da was extracted and automatically integrated using Mass Hunter software. The standards were prepared from a 5 mM CA stock solution in M9 by dilution in methanol, and a calibration curve between 0 and 500 µM CA was obtained (n = 4). Samples were diluted 1:10 in methanol and analyzed within 10 - 15 minutes of storage at RT.

The conversion rate of CA (*Q*_*CA*_, U L^-1^) by tyrosinase was calculated as:

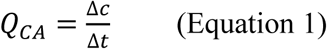

where Δ*c* is the concentration difference (µM) between timepoints and Δ*t* is elapsed time (min). For a tyrosinase concentration of 0.05 mg mL^-1^ an activity of 27.5 ± 3.8 U L^-1^ was determined via this method (Figure S5A).

### Hydrogel bending by mammalian cells

Bilayer hydrogels functionalized with RGD-containing peptides in the responsive layer were prepared and stored overnight in PBS at 4 °C. Prior to cell seeding, hydrogels were equilibrated for 2 h in Dulbecco’s modified Eagle’s medium (DMEM, PAN Biotech, catalog no. P04-03550) supplemented with 10% (v/v) fetal calf serum (FCS, PAN Biotech, catalog no. P30-3602) and 1% (v/v) penicillin-streptomycin solution (P/S, PAN Biotech, catalog no. P06-07100) (referred to as complete medium) under cell culture conditions (37 °C, 5% CO_2_).

Human embryonic kidney cells (HEK-293T, DMSZ ACC 635, RRID: CVCL_0063) were cultured in complete medium at 37 °C in a humid atmosphere with 5% CO_2_. Cells were passaged every 2 – 3 days upon reaching ∼80% confluency. For transient transfection, HEK293T cells (2.3 x 10^5^ cells mL^-1^) were seeded in 10 mL of complete medium per 10 cm dish. After 24 h, cells were transfected using polyethyleneimine (PEI, stock solution: 1 mg mL^-1^ in H_2_O, pH 7, Polyscience, catalog no. 23966-1). For each dish, DNA/PEI complexes were formed by mixing a total of 22.5 µg plasmid DNA (Mock: 21 µg pCDNA3.1 + 1.5 µg plasmid pMB20; PPEP-1: 21 µg pMB5.2 + 1.5 µg pMB20) and 75 µL of PEI stock solution in 1.5 mL of Opti-MEM (Thermo Fisher Scientific, catalog no. 22600-134). After incubation at RT for 20 min, the DNA/PEI complexes were added dropwise to the cells and cells were incubated for 6 h.

Afterwards, HEK-293T (2 x 10^5^ cells mL^-1^) were seeded in 1 mL phenol red–free complete medium (PAN Biotech; cat. no. P04-03591) per 24-well. To initiate hydrogel crosslinking, 10 mg mL^-1^ sterile filtered TG (from 200 mg mL^-1^ in PBS) was added immediately after.

### PPEP-1 activity assays

PPEP-1 activity was determined as follow: 2.5 µg mL^-1^ PPEP-1 was incubated in phosphate buffer (100 mM NaPO_4_, 150 mM NaCl, pH 7) with 25 µM customized Foerster resonance energy transfer (FRET) peptide substrate ((Dabcyl)-KRPNPPVPRK-E(EDANS)-NH_2_, Caslo) at 37°C. Fluorescence was recorded in 2 min intervals for 2 h at 37 °C (excitation: 340 nm, emission: 490 nm). Initial reaction rates were obtained from the slope of the linear fluorescence increase and converted to U L^-^1 using a calibration curve of free EDANS (MedChemExpress, art. no. HY-D1080). Using this method, a specific PPEP-1 activity of 0.012 ± 0.0005 U L^-1^ was determined for 2.5 µg mL^-1^ PPEP-1.

To determine PPEP-1 cell-specific production, mock or PPEP-1 transfected cells were seeded as described above. After 14 and 86 h, medium was collected and stored at −20 °C until further analysis. PPEP-1 activity was measured as described above. Briefly, 50 µL of each sample was incubated with 25 µM FRET peptide at 37 °C and fluorescence (see above) was recorded in 2 min intervals. Initial reaction rates were obtained from the slope of the linear fluorescence increase and converted to enzyme units (U) using an EDANS calibration curve. Cell-specific activity (*Q*_*C*_) was calculated as:

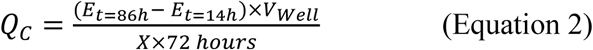

where *E* is the enzymatic activity (U mL^-1^), *V*_*Well*_is the total well volume (1 mL) and *X* is the initial number of transfected cells. Subsequently PPEP-1 specific activity (*Q*_*P*_) was determined as:

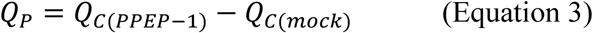

To determine transfection efficiency, cells were detached 14 h post seeding, resuspended in FACS buffer (PBS + 2% FCS), and analyzed via Attune NxT flow cytometer (Thermo Fisher Scientific). Samples were acquired at a flow rate of 200 µL min^-1^ and mVenus fluorescence transferred by the expression vector pMB20 was measured using the BL1 channel (488 nm excitation; 530/30 nm emission). Results were analyzed in FlowJo (v10), and transfection efficiency was calculated as the percentage of mVenus-positive single cells using wild-type HEK-293T cells as the negative control population.

## Statistics

Statistical differences between conditions were assessed by two-way ANOVA and are indicated in the respective experiments. Sample size (n) is indicated in the respective Figure legends and represents replicates. For rheological SAOS measurements, 3 datapoints were obtained for each sample. Data is shown as individual datapoints or mean ± SD.

## Software

Statistical differences between conditions were assessed using GraphPad Prism 10. Figures were partially created with Biorender.com.

During the preparation of the manuscript the authors partly used Microsoft Copilot to improve readability and language. After using this tool, the authors reviewed and edited the content and take full responsibility for the content of the publication.

## Acknowledgements

The authors would like to thank Hanna Mayer for the production of relevant protein. This work was supported by the European Union (ERC, STEADY, 101053857).

## Data Availability Statement

The data that support the findings of this study are available from the corresponding author upon reasonable request.

## Supporting Information

Supporting Information is available from the bioRxiv or from the author.

## Table of contents entry

Here, we present bacterial and mammalian engineered living materials (ELMs) with genetically and enzymatically programmed shape-morphing. This autonomous and reversible process can be triggered by combined endogenous and external stimuli or in a fully autonomous manner by an ELM incorporating out-of-equilibrium counteracting chemical reactions with dynamically changing respective reaction rates.

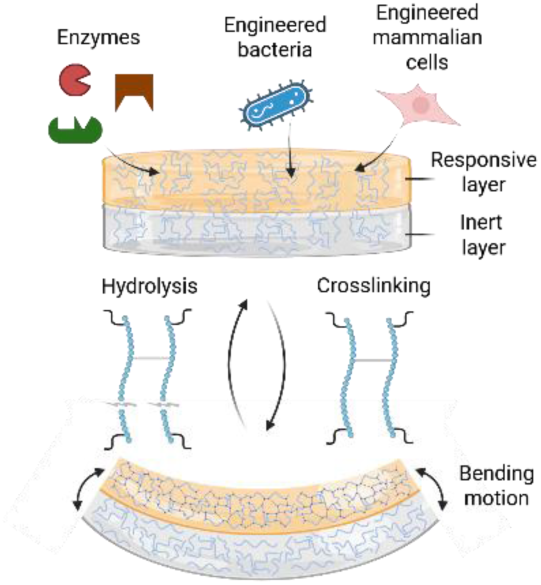

