## Supplementary information for "Genetically Programmed Shape-morphing of Engineered Living Materials"

**Table S1.** Plasmid sequences and annotations.

| Construct | Sequence |
|---|---|
| Tyrosinase<br>(pRSMtyr/<br>pRS39)<br>Backbone<br>T7 promoter<br>Tyrosinase<br>$His_6$-Tag | CCGCGAAATTAATACGACTCACTATAGGGAGACCACAACGGTTTCCCTCTAGAAATAATTTTGTT<br>TAACTTTAAGAAGGAGATATACATATGGGTAACAAATACCGTGTTCGTAAAAACGTACTTCATCT<br>AACAGATACGGAAAAACGTGACTTTGTTCGTACTGTACTTATCCTAAAGGAAAAGGGTATCTACG<br>ATCGTTACATCGCATGGCATGGTGCAGCAGGCAAATTTCATACACCACCAGGCTCTGACCGTAA<br>CGCAGCACATATGTCTTCAGCATTTCTTCCATGGCATCGTGAATACCTTCTACGTTTTGAACGTG<br>ATCTTCAATCAATCAACCCAGAAGTTACACTACCATACTGGGAATGGGAAACGGATGCACAAAT<br>GCAGGACCCATCTCAATCACAGATCTGGTCAGCAGATTTTATGGGTGGCAACGGTAACCCAATC<br>AAAGATTTTATCGTAGACACGGGTCCATTTGCAGCAGGCCGTTGGACTACAATCGACGAACAAG<br>GTAACCCATCAGGTGGCCTTAAACGTAACTTTGGCGCAACAAAAGAAGCACCAACTCTACCAAC<br>ACGTGATGACGTTCTTAACGCACTAAAAATCACGCAGTACGATACTCCACCATGGGACATGACT<br>TCTCAAAACTCATTTCGTAACCAGCTTGAAGGTTTTATCAACGGCCCACAACTACATAACCGTGT<br>TCATCGTTGGGTAGGTGGCCAGATGGGTGTTGTACCAACAGCACCAAACGATCCAGTTTTCTTT<br>CTTCATCATGCAAACGTTGACCGTATCTGGGCAGTATGGCAAATCATCCATCGTAACCAAAACTA<br>CCAGCCAATGAAAAACGGTCCATTTGGCCAGAACTTTCGTGATCCAATGTACCCATGGAACACG<br>ACTCCAGAAGACGTTATGAACCATCGTAAACTTGGCTACGTATACGACATCGAACTACGTAAATC<br>TAAACGTTCTTCACATCATCACCATCACCATTAAAAGCTTGAT |
| PPEP-1<br>(pJB001)<br>Backbone<br>T7 promoter<br>PPEP-1<br>$His_{10}$-Tag | CCGCGAAATTAATACGACTCACTATAGGGGAATTGTGAGCGGATAACAATTCCCCTCTAGAAAT<br>AATTTTGTTTAACTTTAAGAAGGAGATATACCATGGGCCATCATCATCATCATCATCATCATCATC<br>ACAGCAGCGGCCATATCGAAGGTCGTCATATGGATTCAACTACAATACAACAGAATAAAGACAC<br>CCTGTCTCAGATCGTGGTGTTTCCGACCGGTAATTACGATAAAAACGAGGCAAACGCTATGGTT<br>AATCGTCTGGCGAATATTGACGGCAAGTACTTAAACGCGCTGAAACAAAACAATCTGAAGATCA<br>AACTTCTGTCTGGTAAGTTGACGGATGAAAAGGAGTATGCATATCTGAAGGGCGTTGTCCCAAA<br>AGGTTGGGAAGGTACGGGCAAGACCTGGGACGATGTTCCGGGTCTGGGCGGTTCCACCGTGG<br>CTCTGCGCATTGGTTTTAGCAATAAAGGCAAAGGTCATGATGCAATTAACTTGGAACTGCACGA<br>GACTGCGCACGCCATCGACCATATTGTTCTGAACGACATCAGCAAGTCGGCTCAGTTTAAACAA<br>ATCTTTGCGAAAGAGGGCCGTAGCCTGGGAAACGTCAATTACTTGGGGGTGTATCCGGAAGAG<br>TTCTTCGCCGAATCCTTCGCGTACTACTACCTCAACCAGGATACCAACAGCAAATTGAAGTCCG<br>CGTGCCCGCAGACCTATAGCTTCCTGCAAAACTTGGCGAAGTAAGGATCCGGC |
| GyrB -<br>Transformation<br>control<br>(pWW873)<br>Backbone<br>T7 promoter<br>GyrB | CCGCGAAATTAATACGACTCACTATAGGGAGACCACAACGGTTTCCCTCTAGAAATAATTTTGTT<br>TAACTTTAAGAAGGAGATATACATATGTCGAATTCTTATGACTCCTCCAGTATCAAAGTCCTGAAA<br>GGGCTGGATGCGGTGCGTAAGCGCCCGGGTATGTATATCGGCGACACGGATGACGGCACCGG<br>TCTGCACCACATGGTATTCGAGGTGGTAGATAACGCTATCGACGAAGCGCTCGCGGGTCACTG<br>TAAAGAAATTATCGTCACCATTCACGCCGATAACTCTGTCTCTGTACAGGATGACGGGCGCGGC<br>ATTCCGACCGGTATTCACCCGGAAGAGGGCGTATCGGCGGCGGAAGTGATCATGACCGTTCTG<br>CACGCAGGCGGTAAATTTGACGATAACTCCTATAAAGTGTCCGGCGGTCTGCACGGCGTTGGT |

| | |
|---|---|
| His$_6$-Tag | GTTTCGGTAGTAAACGCCCTGTCGCAAAAACTGGAGCTGGTTATCCAGCGCGAGGGTAAAATTC<br>ACCGTCAGATCTACGAACACGGTGTACCGCAGGCCCCGCTGGCGGTTACCGGCGAGACTGAA<br>AAAACCGGCACCATGGTGCGTTTCTGGCCCAGCCTCGAAACCTTCACCAATGTGACCGAGTTC<br>GAATATGAAATTCTGGCGAAACGTCTGCGTGAGTTGTCGTTCCTCAACTCCGGCGTTTCCATTC<br>GTCTGCGCGACAAGCGCGACGGCAAAGAAGACCACTTCCACTATGAAGGCCATCATCACCATC<br>ACCATAAGCTTGATCCGGCTGCTAACAAAGCCCGAAAGGAAGCTGAGTTGGCTGCTGCCACCG<br>CTGAGCAATAACTAGCATAA |
| IL-6-PPEP-1<br>(pMB05.2)<br>Backbone<br>CMV Enhancer +<br>promoter<br>T7 promoter<br>IL-6<br>PPEP-1 | ATACGCGTTGACATTGATTATTGACTAGTTATTAATAGTAATCAATTACGGGGTCATTAGTTCATA<br>GCCCATATATGGAGTTCCGCGTTACATAACTTACGGTAAATGGCCCGCCTGGCTGACCGCCCAA<br>CGACCCCCGCCCATTGACGTCAATAATGACGTATGTTCCCATAGTAACGCCAATAGGGACTTTC<br>CATTGACGTCAATGGGTGGAGTATTTACGGTAAACTGCCCACTTGGCAGTACATCAAGTGTATC<br>ATATGCCAAGTACGCCCCCTATTGACGTCAATGACGGTAAATGGCCCGCCTGGCATTATGCCCA<br>GTACATGACCTTATGGGACTTTCCTACTTGGCAGTACATCTACGTATTAGTCATCGCTATTACCA<br>TGGTGATGCGGTTTTGGCAGTACATCAATGGGCGTGGATAGCGGTTTGACTCACGGGGATTTC<br>CAAGTCTCCACCCCATTGACGTCAATGGGAGTTTGTTTTGGCACCAAAATCAACGGGACTTTCC<br>AAAATGTCGTAACAACTCCGCCCCATTGACGCAAATGGGCGGTAGGCGTGTACGGTGGGAGGT<br>CTATATAAGCAGAGCTCTCTGGCTAACTAGAGAACCCACTGCTTACTGGCTTATCGAAATTAATA<br>CGACTCACTATAGGGAGACCCAAGCTGGCTAGCGTTTAAACTTAAGCTTGGTACCGAGCTCGGA<br>TAGCCACCATGAACTCCTTCTCCACAAGCGCCTTCGGTCCAGTTGCCTTCTCCCTGGGCCTGCT<br>CCTGGTGTTGCCTGCTGCCTTCCCTGCCCCAGATTCTACTACCATCCAACAAAACAAGGACACC<br>TTGTCTCAAATCGTTGTTTTTCCAACTGGTAACTACGATAAGAACGAAGCTAATGCCATGGTTAA<br>CAGATTGGCTAATATCGACGGTAAATACTTGAACGCCTTGAAGCAAAACAACCTGAAGATCAAG<br>TTGTTGTCTGGTAAGTTGACCGACGAAAAAGAATACGCTTACTTGAAAGGTGTTGTCCCAAAAG<br>GTTGGGAAGGTACTGGTAAAACTTGGGATGATGTTCCAGGTTTAGGTGGTTCTACTGTTGCTTT<br>GAGAATTGGCTTTTCCAACAAAGGTAAAGGTCATGATGCCATCAATTTGGAATTGCACGAAACTG<br>CTCATGCCATCGATCATATCGTTTTGAACGATATTTCTAAGAGCGCCCAATTCAAGCAAATCTTC<br>GCAAAAGAAGGTAGATCTTTGGGTAACGTTAACTACTTGGGTGTTTACCCAGAAGAATTTTTCGC<br>TGAATCTTTCGCCTACTACTACTTGAATCAAGACACCAACTCTAAGTTGAAGTCTGCTTGTCCAC<br>AAACCTACTCATTCTTGCAAAATTTGGCTAAATAAGAATTCTGC |
| mVenus<br>(pMB020)<br>Backbone<br>CMV Enhancer +<br>promoter<br>T7 promoter<br>mVenus | ATACGCGTTGACATTGATTATTGACTAGTTATTAATAGTAATCAATTACGGGGTCATTAGTTCATA<br>GCCCATATATGGAGTTCCGCGTTACATAACTTACGGTAAATGGCCCGCCTGGCTGACCGCCCAA<br>CGACCCCCGCCCATTGACGTCAATAATGACGTATGTTCCCATAGTAACGCCAATAGGGACTTTC<br>CATTGACGTCAATGGGTGGAGTATTTACGGTAAACTGCCCACTTGGCAGTACATCAAGTGTATC<br>ATATGCCAAGTACGCCCCCTATTGACGTCAATGACGGTAAATGGCCCGCCTGGCATTATGCCCA<br>GTACATGACCTTATGGGACTTTCCTACTTGGCAGTACATCTACGTATTAGTCATCGCTATTACCA<br>TGGTGATGCGGTTTTGGCAGTACATCAATGGGCGTGGATAGCGGTTTGACTCACGGGGATTTC<br>CAAGTCTCCACCCCATTGACGTCAATGGGAGTTTGTTTTGGCACCAAAATCAACGGGACTTTCC<br>AAAATGTCGTAACAACTCCGCCCCATTGACGCAAATGGGCGGTAGGCGTGTACGGTGGGAGGT<br>CTATATAAGCAGAGCTCTCTGGCTAACTAGAGAACCCACTGCTTACTGGCTTATCGAAATTAATA<br>CGACTCACTATAGGGAGACCCAAGCTGGCTAGCGTTTAAACTTAAGCTTGGTACCGAGCTCGG<br>GCCACCATGGTGAGCAAGGGCGAGGAGCTGTTCACCGGGGTGGTGCCCATCCTGGTCGAGCT<br>GGACGGCGACGTAAACGGCCACAAGTTCAGCGTGTCCGGCGAGGGCGAGGGCGATGCCACCT<br>ACGGCAAGCTGACCCTGAAGTTGATCTGCACCACCGGCAAGCTGCCCGTGCCCTGGCCCACC<br>CTCGTGACCACCCTCGGCTACGGCCTGCAGTGCTTCGCCCGCTACCCCGACCACATGAAGCAG<br>CACGACTTCTTCAAGTCCGCCATGCCCGAAGGCTACGTCCAGGAGCGCACCATCTTCTTCAAG<br>GACGACGGCAACTACAAGACCCGCGCCGAGGTGAAGTTCGATGGCGACACCCTGGTGAACCG<br>CATCGAGCTGAAGGGCATCGACTTCAAGGAGGACGGCAACATCCTGGGGCACAAGCTGGAGTA<br>CAACTACAACAGCCACAACGTCTATATCACCGCCGACAAGCAGAAGAACGGCATCAAGGCAAA<br>CTTCAAGATCCGCCACAACATCGAGGACGGCGGCGTGCAGCTCGCCGACCACTACCAGCAGAA<br>CACCCCCATCGGCGACGGCCCCGTGCTGCTGCCCGACAACCACTACCTGAGCTACCAGTCCAA |

| | |
|---|---|
| | ACTGAGCAAAGACCCCAACGAGAAGCGCGATCACATGGTCCTGCTGGAGTTCGTGACCGCCGC<br>CGGGATCACTCTCGGCATGGACGAGCTGTACAAGGGTAGTGGTAGTGGATCCTGATGAGAATT<br>C |
| Transfection control (pCDNA3.1)<br>Backbone<br>CMV Enhancer + promoter<br>T7 promoter | ATACGCGTTGACATTGATTATTGACTAGTTATTAATAGTAATCAATTACGGGGTCATTAGTTCATA<br>GCCCATATATGGAGTTCCGCGTTACATAACTTACGGTAAATGGCCCGCCTGGCTGACCGCCCAA<br>CGACCCCCGCCCATTGACGTCAATAATGACGTATGTTCCCATAGTAACGCCAATAGGGACTTTC<br>CATTGACGTCAATGGGTGGAGTATTTACGGTAAACTGCCCACTTGGCAGTACATCAAGTGTATC<br>ATATGCCAAGTACGCCCCCTATTGACGTCAATGACGGTAAATGGCCCGCCTGGCATTATGCCCA<br>GTACATGACCTTATGGGACTTTCCTACTTGGCAGTACATCTACGTATTAGTCATCGCTATTACCA<br>TGGTGATGCGGTTTTGGCAGTACATCAATGGGCGTGGATAGCGGTTTGACTCACGGGGATTTC<br>CAAGTCTCCACCCCATTGACGTCAATGGGAGTTTGTTTTGGCACCAAAATCAACGGGACTTTCC<br>AAAATGTCGTAACAACTCCGCCCCATTGACGCAAATGGGCGGTAGGCGTGTACGGTGGGAGGT<br>CTATATAAGCAGAGCTCTCTGGCTAACTAGAGAACCCACTGCTTACTGGCTTATCGAAATTAATA<br>CGACTCACTATAGGGAGACCCAAGCTGGCTAGCGTTTAAACTTAAGCTTGGTACCGAGCTCGGA<br>TCCACTAG |

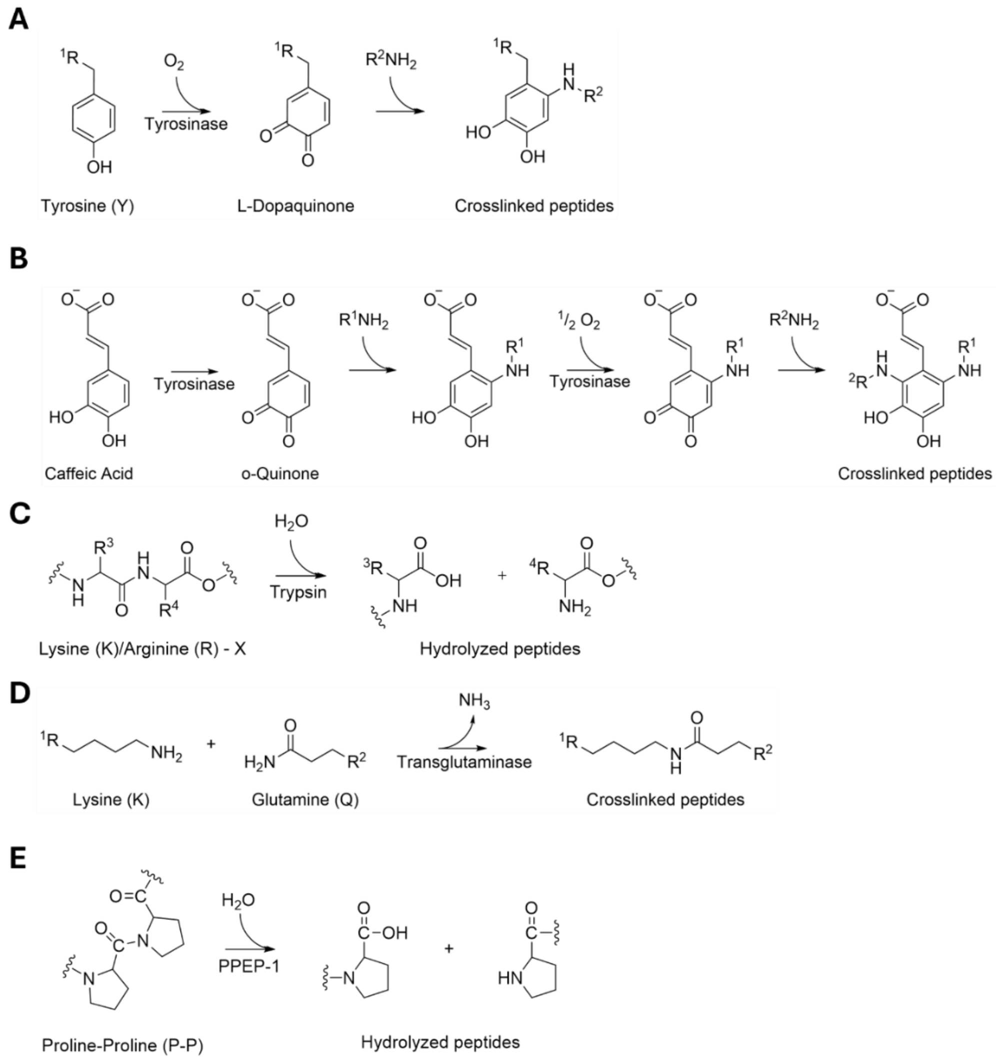


**Figure S1. Chemical reactions catalyzed by enzymes.** (A) Tyrosinase-mediated deswelling via Tyrosine residues. Tyrosine (Y) is oxidized into an L-dopaquinone intermediate by tyrosinase, which subsequently undergoes coupling reactions with Y residues or nucleophilic groups on adjacent peptide chains. (B) Tyrosinase-mediated deswelling via caffeic acid (CA). CA is converted by tyrosinase into a reactive o-quinone intermediate which performs a nucleophilic attack of lysine ε-amino groups, forming covalent crosslinks between two peptides. (C) Trypsin-mediated hydrolysis. Peptide bonds at the C-terminus of lysine (K) or arginine (R) residues are hydrolysed by trypsin. (D) Transglutaminase (TG) -mediated deswelling. TG catalyses an acyl-transfer reaction between the γ-carboxamide group of glutamine (Q) residues and the ε-amino group of K residues, forming covalent isopeptide bonds that crosslink peptide chains. (E) PPEP-1-mediated hydrolysis. PPEP-1 catalyses the hydrolysis of a proline – proline bond within the target sequence PNP↓PVP.

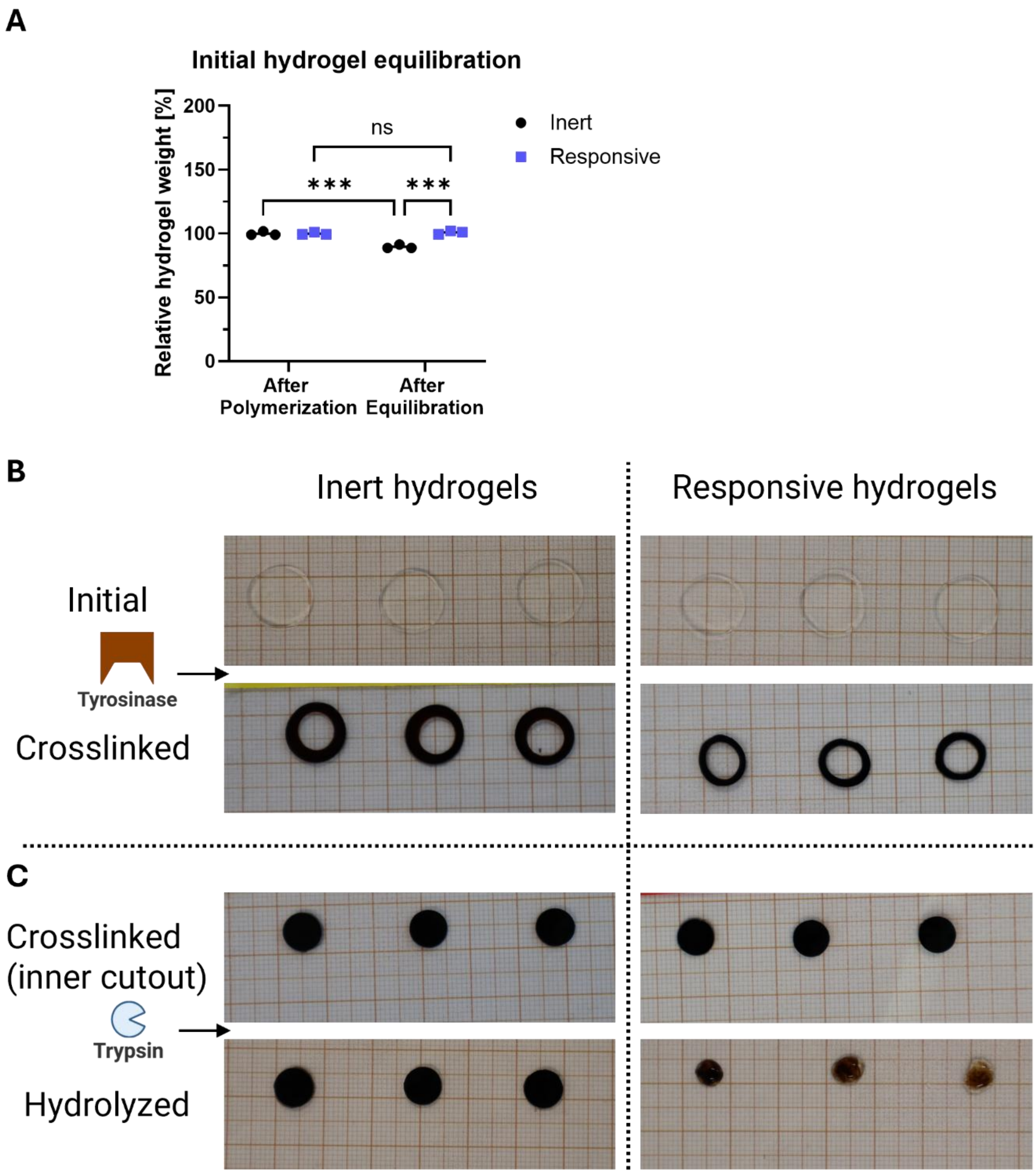


**Figure S2. Initial swelling and size changes of inert and responsive hydrogels upon tyrosinase or trypsin treatment.** (A) Initial hydrogel equilibration in M9. Inert and responsive hydrogel weight differences after polymerization and after equilibration in M9 were assessed. Statistical significance of replicates (n = 3) between conditions was assessed by two-way ANOVA; ns = not significant, * = $p \leq 0.033$, ** = $p \leq 0.002$, *** = $p \leq 0.001$. (B + C) Imaging of enzyme-mediated hydrogel changes. Supplementary information to Figure 2. Initial images were taken after cutting the hydrogel from (A) with a biopsy punch to 12 mm diameter. After tyrosinase-mediated deswelling (B), 8 mm diameter hydrogels were cut out and treated with trypsin (C).

**Table S2.** Absolute values for hydrogel weight at different stages of processing for tyrosinase and trypsin treatment. After swelling hydrogels were cut to 12 mm diameter providing the initial value; after deswelling hydrogels were cut to 8 mm diameter providing the crosslinked (inner cutout) value.

| **Hydrogel state** | **Inert hydrogels [mg]** | **Responsive hydrogels [mg]** |
|---|---|---|
| After polymerization | 116.0 ± 1.7 | 115.7 ± 1.1 |
| After equilibration | 104.0 ± 1.7 | 116.7 ± 1.5 |
| Initial | 104.0 ± 1.7 | 110.0 ± 6.1 |
| Crosslinked | 92.7 ± 4.5 | 46.8 ± 7.7 |
| Crosslinked (inner cutout) | 41.0 ± 1.7 | 32.3 ± 1.2 |
| Hydrolyzed | 43.7 ± 3.5 | 5.0 ± 1.7 |

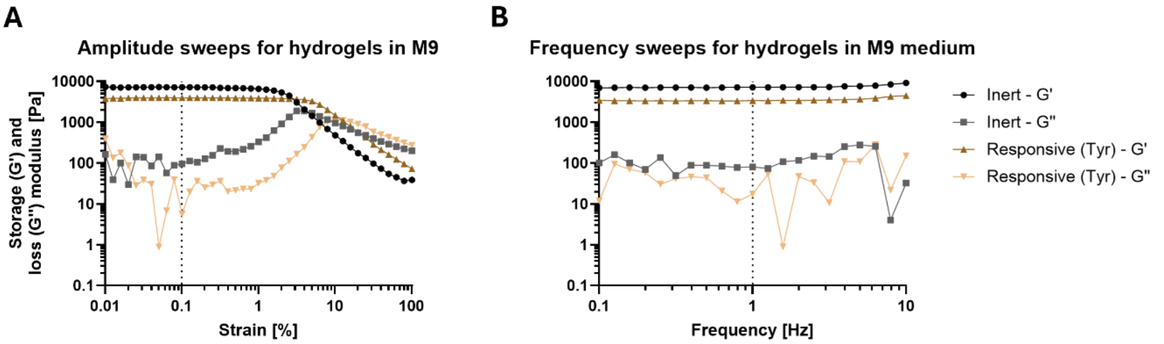


**Figure S3. Rheological characterization of tyrosinase- and trypsin-responsive hydrogels.** (A + B) Rheological amplitude (A) and frequency (B) sweeps. Storage (G') and loss (G'') modulus measurements were performed with inert and responsive hydrogels equilibrated in M9 to determine the linear viscoelastic (LVE) range. Dotted lines indicate the parameters selected for further analysis.

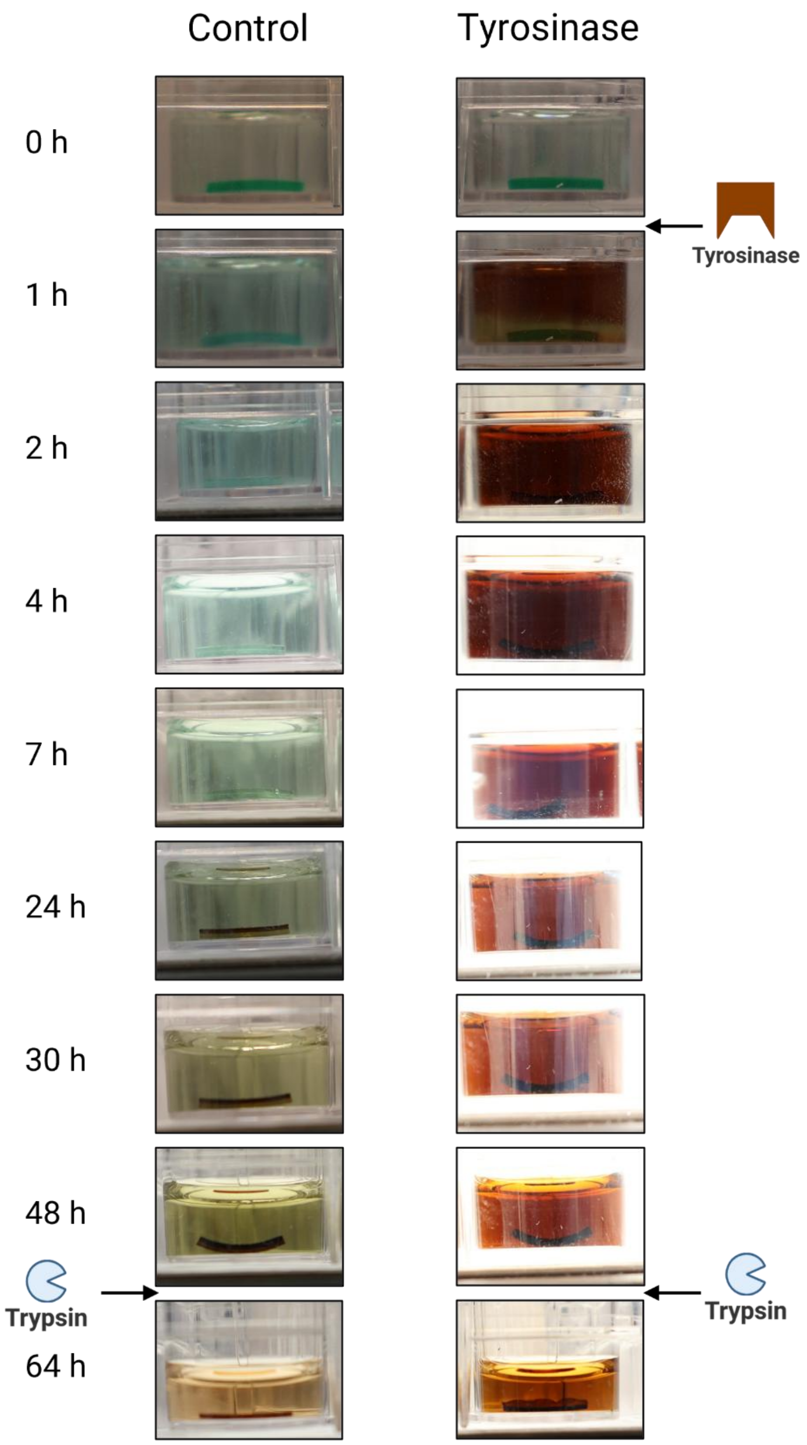


**Figure S4. Bilayer hydrogel bending upon tyrosinase or trypsin treatment.** Supplementary information to Figure 3 showing original images from tyrosinase- (0 - 48 h) and trypsin- (48 - 64 h) mediated hydrogel bending after the indicated incubation time. Control hydrogels were placed in M9 containing CA but no enzyme. To improve visual clarity and ensure object focus, exposure time was adjusted for tyrosinase-treated samples using the built-in settings of the digital camera. Depicted images are representative for replicates (n = 3).

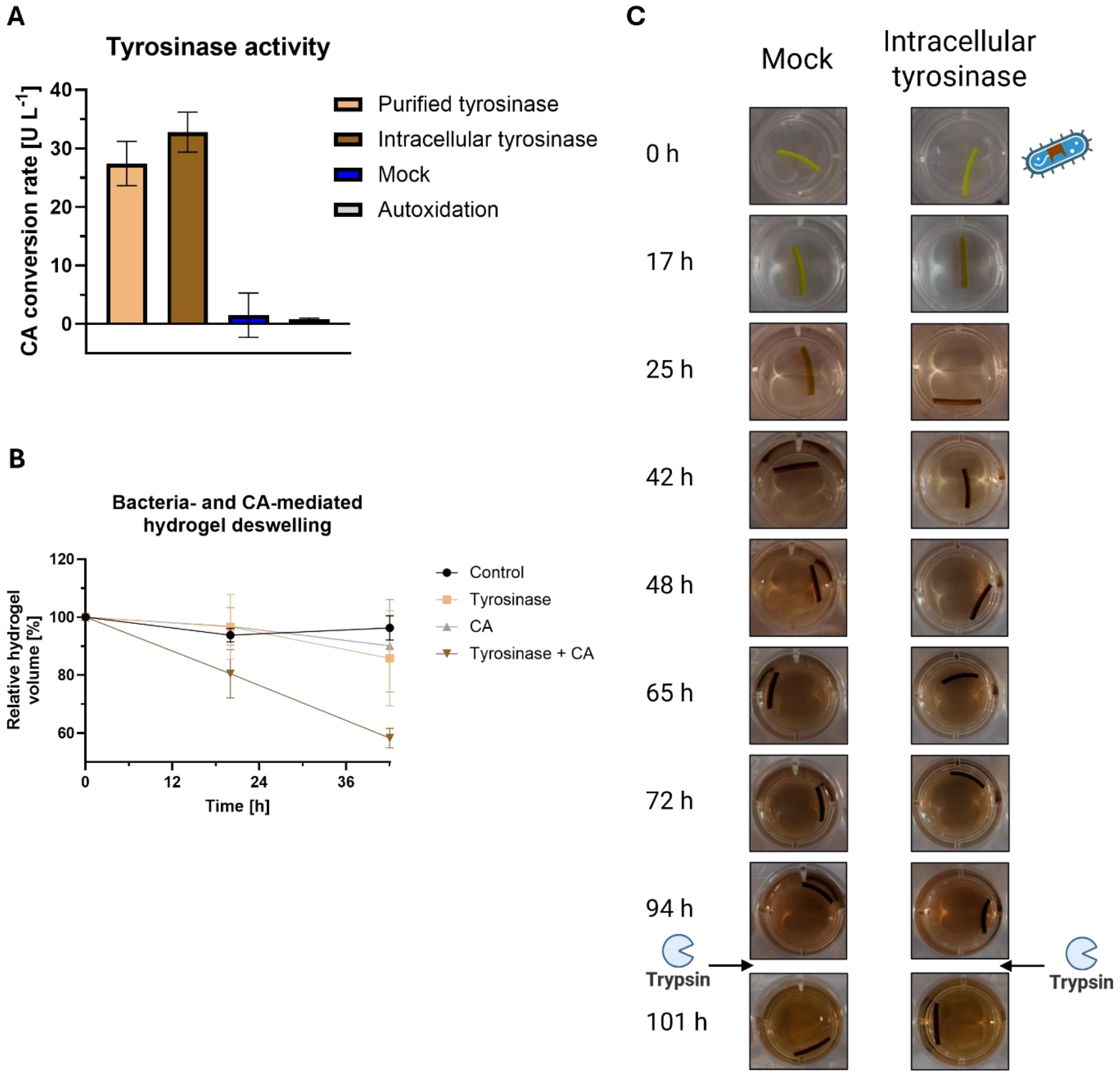


**Figure S5. Hydrogel bending by embedded bacteria producing intracellular tyrosinase.** (A) Enzyme-mediated conversion and autoxidation of CA. Engineered bacteria producing intracellular tyrosinase or GyrB (mock), as well as purified tyrosinase (50 µg mL$^{-1}$) were incubated in M9 containing CA (5 mM) for 60 or 90 min. As control, CA was incubated in the absence of enzyme for 60 or 1500 min. CA concentration difference between the time points was analyzed via Liquid Chromatography coupled to Electrospray Ionization Time-Of-Flight Mass Spectrometry (LC-ESI-QTOF-MS) and enzymatic activity calculated according to Equation 1. (B) Bacteria- and CA-mediated hydrogel deswelling. Responsive hydrogels containing fluorescent RGD peptide and embedded bacteria were prepared and equilibrated for 3 h in M9 containing isopropyl-$\beta$-D-thiogalactopyranoside (IPTG) at 30 °C and 100 rpm. Afterward, CA was added to the respective samples and incubation continued for 42 h. At 16 h, medium was replaced with fresh M9 supplemented with IPTG and CA. Images were taken before (0 h) and after CA addition at the indicated timepoints and volumetric deswelling was calculated from hydrogel diameter assuming isotropic behavior. (C) Imaging of bacteria-mediated hydrogel deformation. Supplementary information to Figure 4 showing original images from bacteria-driven hydrogel bending via intracellular tyrosinase (0 - 94 h), as well as trypsin driven reversion of deformation (94 - 101 h).

Depicted images are representative for replicates (n = 3); values are shown as mean ± SD.

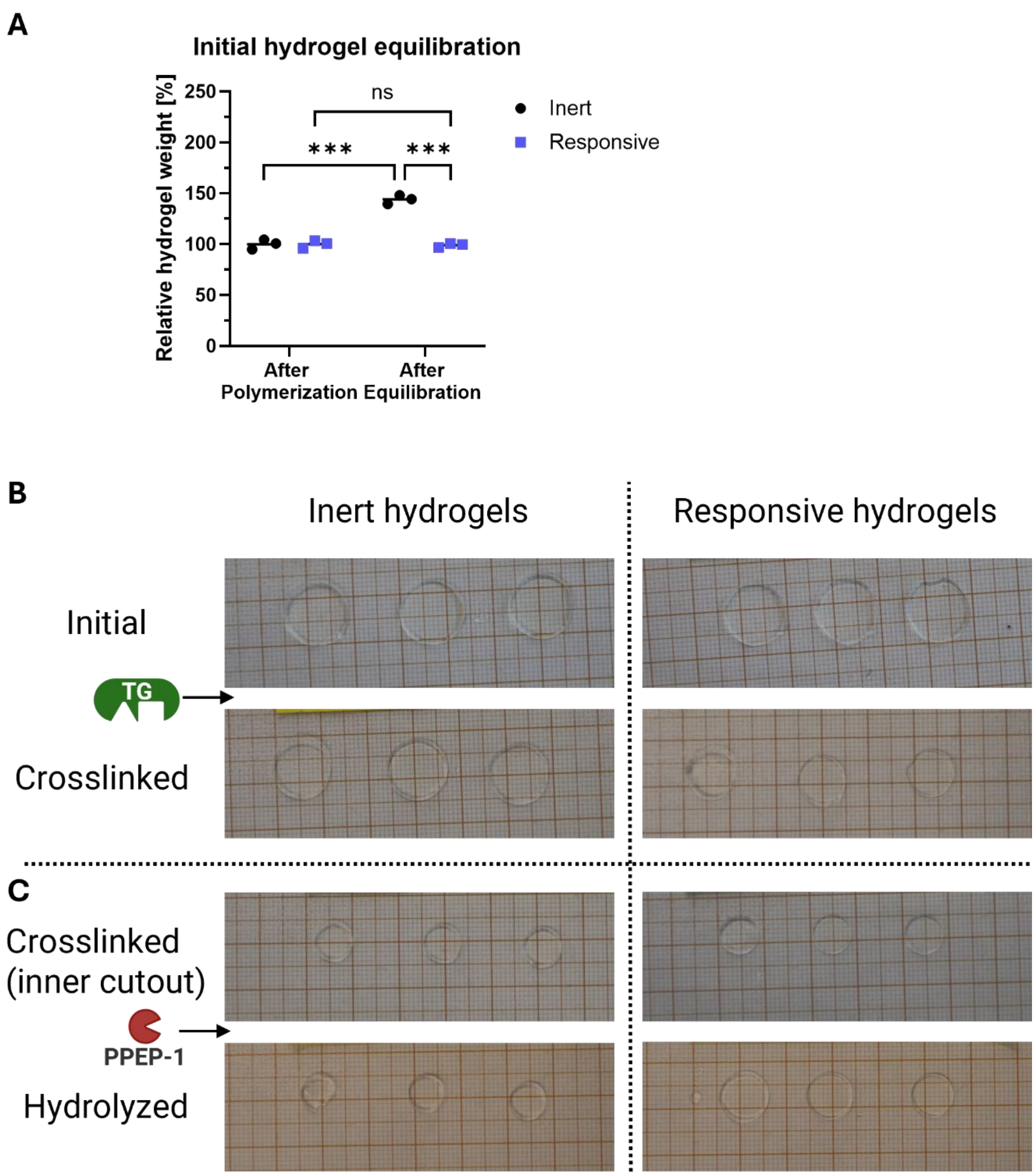


**Figure S6. Initial swelling and size changes of inert and responsive hydrogels upon transglutaminase or PPEP-1 treatment.** (A) Initial hydrogel equilibration in PBS. Inert and responsive hydrogel weight differences after polymerization and after equilibration in PBS were assessed. (B - C) Imaging of enzyme-mediated hydrogel changes. Supplementary information to Figure 5. Initial images were taken after cutting hydrogels from (A) to 12 mm diameter. After TG-mediated deswelling (B), 8 mm diameter hydrogels were cut out and treated with PPEP-1 (C).

Statistical significance of replicates (n = 3) between conditions was assessed by two-way ANOVA; ns = not significant, * = $p \leq 0.033$, ** = $p \leq 0.002$, *** = $p \leq 0.001$.

**Table S3.** Absolute values for hydrogel weight at different stages of processing for TG and PPEP-1 treatment. After swelling hydrogels were cut to 12 mm diameter providing the initial value; after deswelling hydrogels were cut to 8 mm diameter providing the crosslinked (inner cutout) value.

| **Hydrogel state** | **Inert hydrogels [mg]** | **Responsive hydrogels [mg]** |
|---|---|---|
| After polymerization | 105.3 ± 5.0 | 105.3 ± 4.0 |
| After equilibration | 151.7 ± 4.5 | 104.3 ± 2.1 |
| Initial | 124.0 ± 8.9 | 89.0 ± 8.7 |
| Crosslinked | 121.3 ± 2.3 | 52.3 ± 4.5 |
| Crosslinked (inner cutout) | 48.0 ± 1.4 | 31.0 ± 2.7 |
| Hydrolyzed | 44.0 ± 2.8 | 56.7 ± 7.4 |

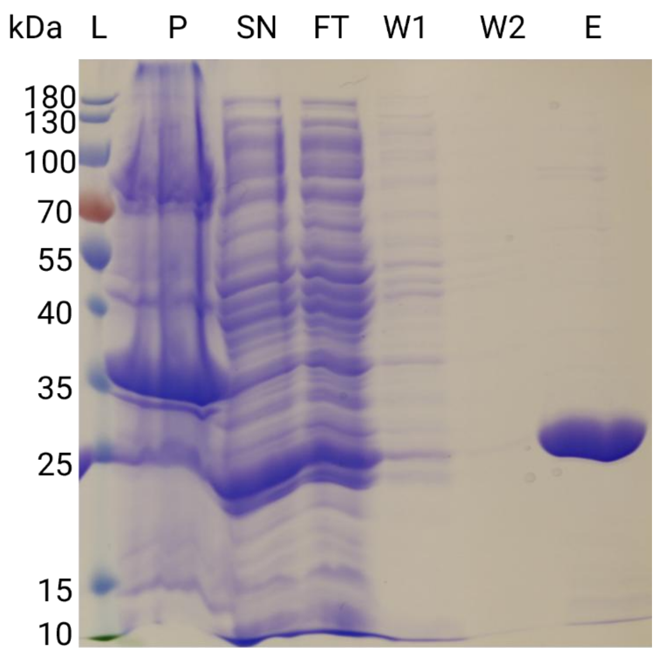


**Figure S7. Characterization of recombinantly purified PPEP-1.** Fractions of purification process by IMAC were collected and visualized via SDS-PAGE followed by Coomassie staining. (L: molecular ladder, P: insoluble fraction of *E. coli* lysate; S: soluble fraction of lysate; F: flow-through of IMAC column; W1/W2: wash of column; E: eluate of column). Expected molecular weight: 24.2 kDa.

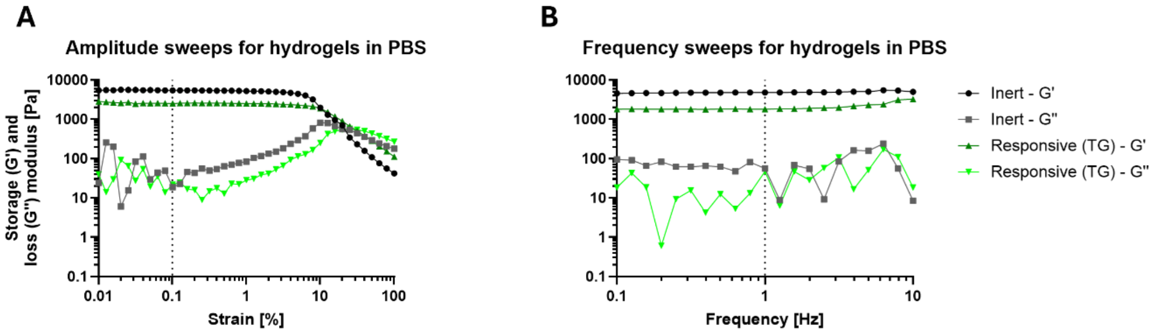


**Figure S8. Initial swelling and rheological characterization of transglutaminase and PPEP-1 responsive hydrogels.** (A + B) Rheological amplitude (A) and frequency (B) sweeps. G’ and G’’ of inert and responsive hydrogels equilibrated in PBS were measured to determine the linear viscoelastic (LVE) range. Dotted lines indicate the parameters selected for further analysis.

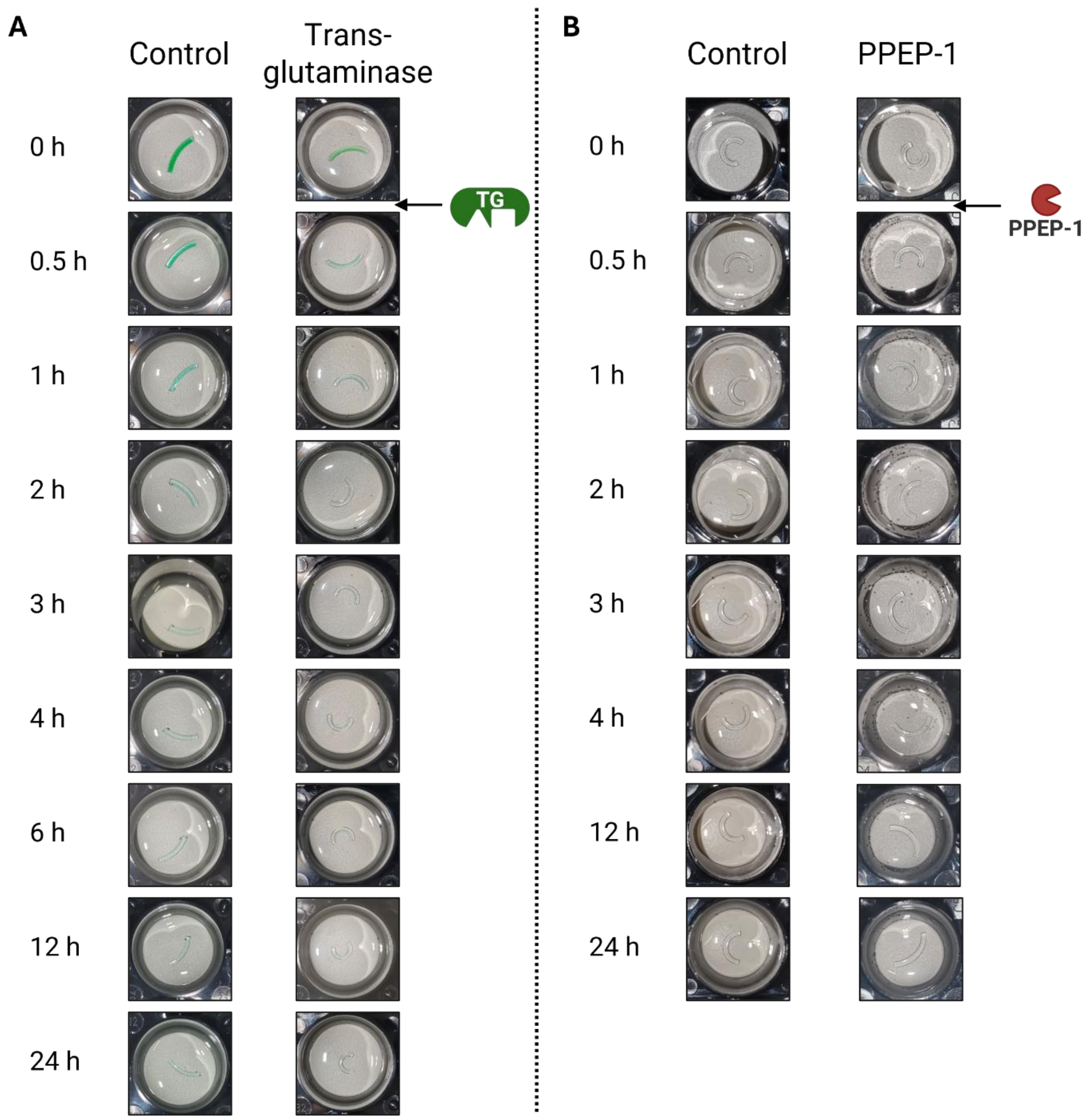


**Figure S9. Bilayer hydrogel bending upon transglutaminase and PPEP-1 treatment.** (A + B) Supplementary information to Figure 6 showing original images from TG-mediated hydrogel bending (A) as well as PPEP-1 driven reversion of deformation (B) after the indicated incubation times. Control hydrogels were placed in enzyme-free PBS. Depicted images are representative for replicates (n = 3).

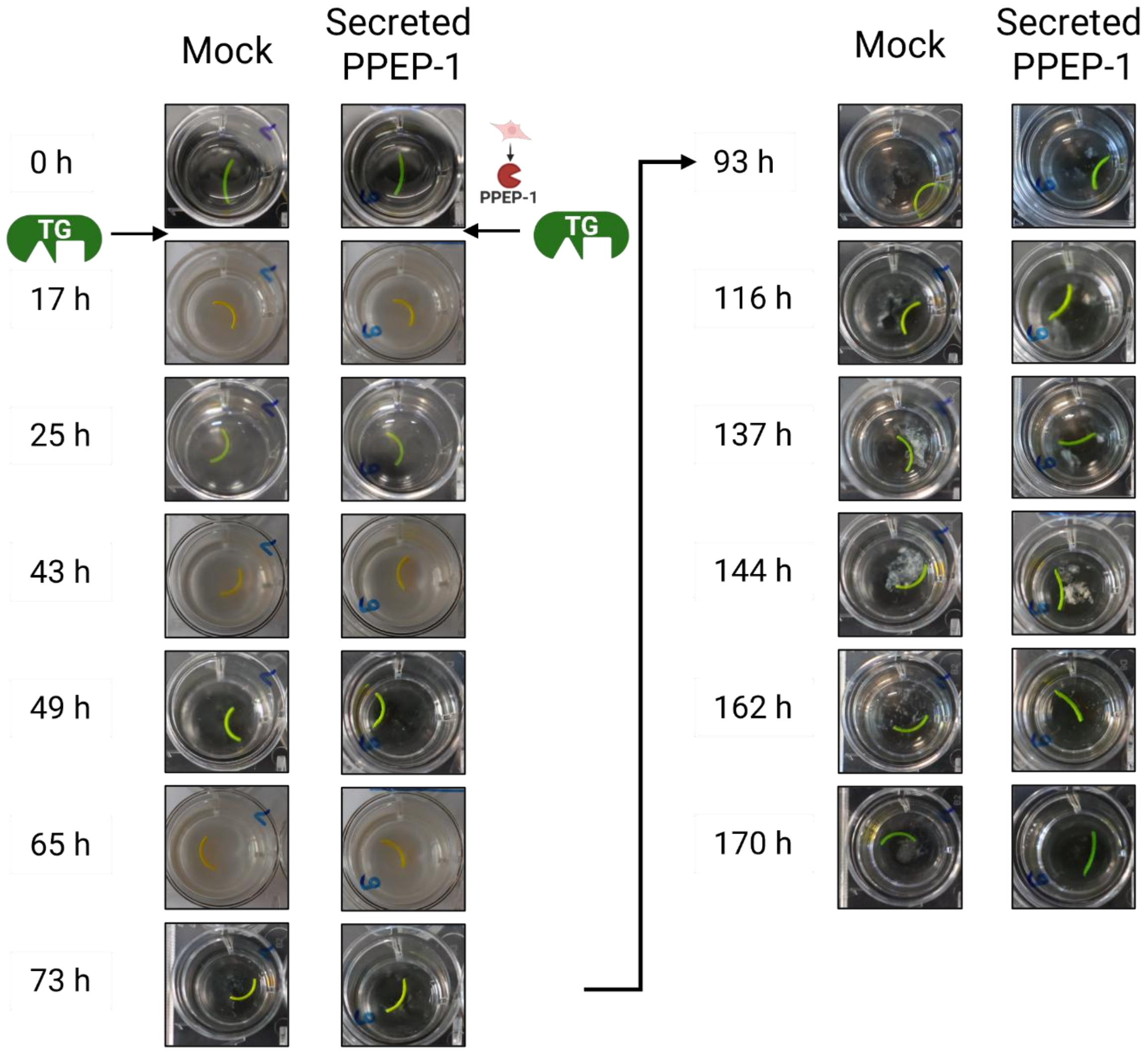


**Figure S10. Hydrogel bending by mammalian cells secreting PPEP-1.** Supplementary information to Figure 7 showing original images of mammalian cell driven hydrogel bending via cell secreted PPEP-1 after TG treatment at the indicated time points. Depicted images are representative for replicates (n = 3).
